# High-Resolution Spatiotemporal Analysis of Single Serotonergic Axons in an *In Vitro* System

**DOI:** 10.1101/2022.07.16.500161

**Authors:** Melissa Hingorani, Adele M.L. Viviani, Jenna E. Sanfilippo, Skirmantas Janušonis

## Abstract

Vertebrate brains have a dual structure in that they are composed of (*i*) axons that can be well captured with graph-theoretical methods and (*ii*) axons that form a dense matrix in which neurons with precise connections operate. A core part of this matrix is formed by axons (fibers) that store and release 5-hydroxytryptamine (5-HT, serotonin), an ancient neurotransmitter that supports neuroplasticity and has profound implications for mental health. The self-organization of the serotonergic matrix is not well understood, despite recent advances in experimental and theoretical approaches. In particular, individual serotonergic axons produce highly stochastic trajectories, fundamental to the construction of regional fiber densities, but further advances in predictive computer simulations require more accurate experimental information. This study examined single serotonergic axons in culture systems (co-cultures and monolayers), by using a complementary set of high-resolution methods: confocal microscopy, holotomography (refractive index-based live imaging), and super-resolution (STED) microscopy. It shows that serotonergic axon walks in neural tissue may strongly reflect the stochastic geometry of this tissue and it also provides new insights into the morphology and branching properties of serotonergic axons. The proposed experimental platform can support next-generation analyses of the serotonergic matrix, including seamless integration with supercomputing approaches.

## 1. INTRODUCTION

Virtually all neural networks in vertebrate brains operate inside a dense matrix of axons whose individual trajectories are inherently unpredictable. Network connections, as well as connections among neuron types, can be well described with graph-theoretical approaches which are strongly deterministic and topological (Sporns et al., 2005; Jiang et al., 2015; Lynn and Bassett, 2019; Faskowitz et al., 2022). In contrast, descriptions of the axon matrix require methods that are inherently stochastic (probabilistic) and geometric (distance-respecting) (Janušonis et al., 2019; Janušonis et al., 2020). Therefore, the brain can be viewed as “a graph embedded in a cotton ball,” comprising strongly “deterministic” and strongly “stochastic” axons. This duality may be a manifestation of the ubiquitous but subtle balance between determinism and chaos, necessary to achieve self-organization and complexity in nature (Teuscher, 2022). It may also be mimicked in a primitive form in the “dropout” technique of artificial neural networks, in which better regularization is achieved by knocking out a random subset of “neurons” in each training iteration, by a process external to the network (Goodfellow et al., 2016; Labach et al., 2019).

Neuroanatomically, the axons that form this matrix are known as the “ascending reticular activating system” (ARAS), due to the fact that they primarily (but not exclusively) originate in the brainstem. A large class of these axons contain and release 5-hydroxytryptamine (5-HT, serotonin), a neurotransmitter that dates back to the origin of all *Bilateria* (Moroz et al., 2021). They are classically referred to as “serotonergic axons,” but currently they are more accurately interpreted as axons of a heterogeneous group of neurons that cluster in specific brain regions and are unique in their expression of tryptophan hydroxylase 2 (*Tph2*), a rate-limiting enzyme in the 5-HT synthesis pathway (Ren et al., 2018; Okaty et al., 2019). In mammals, all serotonergic neurons are restricted to the brainstem, within rather loosely organized raphe nuclei (Jacobs and Azmitia, 1992; Hornung, 2003). The raphe region has been strongly conserved in evolution (Butler and Hodos, 2005). However, in cartilaginous fish, serotonergic neurons can also be present in the hypothalamus, especially around the infundibulum, as well as in some other brain regions (Stuesse et al., 1991; Carrera et al., 2008). In cartilaginous and bony fish, serotonergic neurons can appear in the habenula during development (Ebbesson et al., 1992; Carrera et al., 2008). In the fish and reptilian spinal cord, non-serotonergic neurons can become serotonergic in response to injury (Fabbiani et al., 2018; Huang et al., 2021), and intraspinal serotonergic neurons have also been reported in mammals (Huang et al., 2021). It can therefore be hypothesized that the raphe region is consistently conducive to this phenotype, but that it can also be induced elsewhere in the central nervous system.

In mammalian brains, serotonergic neurons are among the earliest neurons to mature: 5-HT synthesis has been detected around embryonic days 11-13 in mice and rats (Lidov and Molliver, 1982; Hendricks et al., 1999; Hawthorne et al., 2010) and around 5 weeks of gestation in humans (Sundstrom et al., 1993; Mai and Ashwell, 2004). In further development, they extend axons that initially travel along well-defined paths but eventually disperse in the entire brain in a diffusion-like process (Lidov and Molliver, 1982; Aitken and Tork, 1988; Kiyasova and Gaspar, 2011; Jin et al., 2016; Maddaloni et al., 2018; Donovan et al., 2019). These axons grow to become extremely long; because the proximal and distal ends of a given segment cannot be readily identified outside the brainstem, serotonergic axons are often called “fibers.” According to early estimates, each cortical neuron of the rat brain may receive around 200 serotonergic varicosities (putative dilated axon segments) (Jacobs and Azmitia, 1992), which is generally consistent with the current notion that serotonergic axons contribute to pericellular baskets (Senft and Dymecki, 2021).

The developmental axon dispersal eventually leads to the formation of the serotonergic matrix which varies in density across brain regions. Over the past decades, these densities have been mapped in detail in several mammalian species (Steinbusch, 1981; Foote and Morrison, 1984; Lavoie and Parent, 1991; Vertes, 1991; Voigt and de Lima, 1991; Morin and Meyer-Bernstein, 1999; Vertes et al., 1999; Way et al., 2007; Linley et al., 2013). In humans, altered serotonergic densities have been associated with various mental disorders and conditions: Autism Spectrum Disorder (Azmitia et al., 2011), epilepsy (Maia et al., 2019), Major Depressive Disorder (Numasawa et al., 2017), and exposure to 3,4-methylenedioxy-methamphetamine (MDMA, Ecstasy) (Adori et al., 2011). Generally, serotonergic signaling is thought to promote plasticity (Lesch and Waider, 2012; Campanelli et al., 2021), perhaps by keeping synapses in a functionally semi-solidified state. It may explain the profound effects of 5-HT-like psychedelics in mental disorders (Vollenweider and Kometer, 2010; Daws et al., 2022). A growing body of literature suggests close associations between plasticity and randomness in other systems (van der Groen et al., 2022; Voortman and Johnston, 2022), suggesting that the stochasticity of serotonergic axons may not be an evolutionary accident.

The self-organization of the brain serotonergic matrix is a fundamental problem that remains unsolved. Effectively, one has to find a process that starts with the strongly stochastic trajectories of individual serotonergic axons (unique in each individual at the microscopic level) and ends with the regionally-specific serotonergic densities (consistent across individuals at the macroscopic level). It is natural to assume that this process is controlled by molecular signals. Experimental research suggests that they can include the LIM homeodomain protein *Lmx1b* (Donovan et al., 2019), protocadherin α (*Pcdh-αC2*) (Katori et al., 2009; Chen et al., 2017; Katori et al., 2017), the growth factor S100β (Whitaker-Azmitia, 2001), the brain-derived neurotrophic factor (BDNF) (Mamounas et al., 1995), and, importantly, 5-HT itself (Migliarini et al., 2013; Nazzi et al., 2019). The significance of 5-HT as a signal is in that it creates a simple feedback loop, capable of regulating axon growth independently of highly orchestrated developmental sequences (analogously to the development of microvasculature controlled by hypoxia-inducible factor 1α). Notably, serotonergic axons are nearly unique in their ability to regenerate in the adult mammalian brain (Jin et al., 2016; Kajstura et al., 2018; Cooke et al., 2022). This potential suggests that individual axons may get routinely interrupted in normal neural tissue, which is likely considering their typically small diameter, extreme trajectory lengths, various tension forces (*e*.*g*., some serotonergic axons travel through densely packed white matter), and glial activity (Šmít et al., 2017; Janušonis et al., 2019; Janušonis et al., 2020). It further suggests that serotonergic densities are not only built in development but may also have to be actively maintained in the adult brain.

In addition to these biologically-motivated approaches, regionally-specific densities can arise as a consequence of the stochastic properties of serotonergic axons, including their interaction with boundaries and obstacles (Janušonis and Detering, 2019; Janušonis et al., 2019; Janušonis et al., 2020; Vojta et al., 2020). This interdisciplinary approach capitalizes on recent advances in applied mathematics, such as reflected fractional Brownian motion (Wada and Vojta, 2018), and can be readily implemented in supercomputing simulations that bridge the micro- and macro-scales. It has demonstrated that serotonergic densities may depend on the flexibility of individual axons (Janušonis and Detering, 2019), as well as on the curvature of tissue boundaries, with intriguing implications for comparative and clinical neuroscience.

Further progress in this research requires a deeper understanding of single serotonergic axons. Studies in brain tissue are technically challenging because of their high density, small caliber (which approaches the limit of optical resolution), and trajectories that continuously change their orientation. In particular, axons routinely cross at submicrometer distances, making their individual discrimination difficult or impossible (Janušonis et al., 2019). Several groups, including ours, have attempted to analyze individual serotonergic axons in fixed brain tissue (Gagnon and Parent, 2014; Maddaloni et al., 2017; Janušonis and Detering, 2019; Janušonis et al., 2019). However, this information remains insufficient for reliable modeling. The important unsolved problems include the extent to which serotonergic axons reflect the stochastic geometry of the surrounding neural tissue, the interpretation of varicosity-like profiles (in terms of growth dynamics), the stability of other morphological features along the trajectory, branching patterns and frequency, and others.

Individual axons can be made easily accessible in primary neuronal cell cultures, at the expense of environments that may not be natural in their dimensionality, packing, viscoelasticity, and other characteristics. Nevertheless, cell cultures are well positioned to reveal fundamental characteristics of axons, such as extension, varicosity formation, or fasciculation (Šmít et al., 2017; Yurchenko et al., 2019; Sun et al., 2022). These local properties can then be computationally extended to longer times and distances, with further experimental verifications (including normal brain tissue).

Serotonergic neuron cultures were pioneered several decades ago and have been used in morphological, electrophysiological, and pharmacological studies (Lauder et al., 1982; Azmitia and Whitaker-Azmitia, 1987; Johnson, 1994; Johnson and Yee, 1995; Wang et al., 1998; Lautenschlager et al., 2000; Nishi et al., 2000; Yasufuku-Takano et al., 2008; Scheuch et al., 2010; Montgomery et al., 2014; Mercer et al., 2017). Virtually all of these studies focused on population-level descriptions. The present study is the first high-resolution analysis of single serotonergic axons *in vitro* that integrates information obtained with 3D-confocal imaging, 4D-holotomography, and super-resolution microscopy. It demonstrates the potential of these approaches in further experimental studies and immediately informs modeling efforts.

## 2. METHODS

All procedures have been approved by the UCSB Institutional Animal Care and Use Committee.

### 2.1. Primary co-cultures of midbrain neurons and cortical glia

The procedures were based on the protocol developed in Dr. David Sulzer’s laboratory (Columbia University) (Staal et al., 2007).

In the first step, a monolayer of glial cells (primarily astrocytes) was produced. Rat pups (Sprague-Dawley, Charles River, postnatal days (PD) 1-3) were anesthetized on ice, decapitated, and their cerebral cortex was dissected under a stereoscope with fine surgical tools. The collected tissue (from around 2 pups) was placed in Dulbecco’s phosphate buffered saline (DPBS; Sigma-Aldrich # D1408) on ice and cut into small (around 1 mm^3^) pieces. The pieces were immediately transferred into a glia-specific papain solution effused with carbogen (95% O_2_ and 5% CO_2_) to dissociate the cells. The papain solution was composed of papain (20 Units/mL; Worthington Biochemical Corporation #LS003126), 1 mM cysteine (from cysteine water, described below), 1× H&B concentrate (described below), and 0.001% phenol red (all concentrations are final). The cysteine water contained 1.25 mM L-cysteine and 1.9 mM CaCl_2_. The 5× H&B concentrate contained 116 mM NaCl, 5.4 mM KCl, 26 mM NaHCO_3_, 2 mM NaH_2_PO_2_·H_2_O, 1 mM MgSO_4_, 0.5 mM EDTA, and 25 mM glucose. Following cell dissociation, cells were washed, gently triturated with GSM (described below), and counted. Cells were diluted to a density of 1,000,000-1,500,000 cells/mL and plated at around 80,000 cells per culture dish. The 35 mm-culture dishes with a bottom glass coverslip (No. 1.5) were pre-coated with poly-D-lysine (Mattek #P35GC-1.5-14-C) and further coated with laminin (at 10 µg/mL; Sigma-Aldrich #CC095). The glia-specific medium (GSM) was composed of Minimum Essential Medium Eagle (MEM) (180 mL; Sigma-Aldrich #M2279), fetal bovine serum (not heat-inactivated, 20 mL; ThermoFisher # 26140087), glucose (1.5 mL of a 45% solution; Sigma-Aldrich # G8769), insulin (40 µL of 25 mg/mL [0.02 M HCl]; Sigma-Aldrich #I5500), glutamine (0.5 mL of a 200 mM solution; Sigma-Aldrich #G2150), and penicillin-streptomycin (0.24 mL of a solution containing 10,000 Units/mL penicillin and 10 mg/mL streptomycin; Sigma-Aldrich #P0781). When the glia (feeder) layer became 70% confluent (3-5 days after plating), 5-fluoro-2’-deoxyuridine (FDU) was added to inhibit non-neuronal cell proliferation. The FDU stock solution was prepared by adding 15 mL of a uridine solution (16.5 mg/mL; Sigma-Aldrich #U3003) to 100 mg of FDU (FDU; Sigma-Aldrich # F0503). Before use, it was diluted by adding 0.2 mL of the stock to 1.8 mL MEM, and 20 µL of the diluted solution was added to each dish with 2 mL of the glia-specific medium.

In the second step (around 7-14 days after the initial plating), midbrain neurons were added to the culture. Mouse pups (C57BL/6, Charles River, PD 1-2) were anesthetized on ice, decapitated, and their midbrain at the level of the rostral raphe nuclei was dissected under a stereoscope with fine surgical tools. The collected tissue (from 5-7 pups) was placed in PBS on ice and cut into small (around 1 mm^3^) pieces. The pieces were immediately transferred into a neuron-specific papain solution effused with carbogen to dissociate the cells. The papain solution was composed of papain (20 Units/mL), 1 mM L-cysteine (from cysteine water), 1× H&B concentrate, 3.75 mN HCl, 0.5 mM kynurenic acid (from a 0.5 M solution [in 1 N NaOH]; Sigma-Aldrich #K3375), and phenol red (0.001%) (all concentrations are final). Following cell dissociation, cells were washed, triturated with cNSM (described below), and counted. Cells were diluted and to a density of 1,000,000 cells/mL and plated at around 60,000-80,000 cells per dish. Cells were plated in slide rings (Thomas Scientific # 6705R12) on glia monolayers to ensure neurons adhere to the coverslips and do not get washed away. The midbrain neuron-specific medium (cNSM) was composed of MEM (94 mL), Dulbecco’s Modified Eagle’s Medium (low glucose) (80 mL; Sigma-Aldrich #D5546), heat-inactivated fetal bovine serum (2 mL; ThermoFisher #A3840301), glucose (1.5 mL of a 45% solution), glutamine (0.5 mL of a 200 mM solution), bovine serum albumin (fraction V) (0.5 g; Sigma-Aldrich # A4503), Ham’s F-12 nutrient mixture (20 mL; Sigma-Aldrich # N4888), catalase in an aqueous solution (0.1 mL; Sigma-Aldrich # C3155), kynurenic acid (200 µL of a 0.5 M solution), HCl (50 µL of a 5 N solution), and the di Porzio concentrate (2 mL). The di Porzio concentrate (di Porzio et al., 1980; Casper et al., 1991) was composed of 6.25 µg/mL progesterone (Sigma-Aldrich #P0130), 4 µg/mL corticosterone (Sigma-Aldrich #C2505), 2.5 mg/mL insulin, 0.52 µg/mL Na_2_SeO_3_ (Sigma-Aldrich #214485), 2 µg/mL 3,3’,5-triiodo-L-thyronine sodium salt (Sigma-Aldrich #T2752), 0.5 mg/mL superoxide dismutase (Sigma-Aldrich #S7571), 0.24 mg/mL putrescine dihydrochloride (Sigma-Aldrich #P7505), and 10 mg/mL apo-transferrin (Sigma-Aldrich #T1428) in Hanks’ Balanced Salt Solution (HBSS) (ThermoFisher #14170120). The neuron-specific medium was additionally pre-conditioned for 24 hours in either confluent glia cultures (in the original culture dishes) or T225 flasks containing glial cell monolayers. Two hours after plating, the slide rings were removed and glial-derived neurotrophic factor (GDNF) was added at the final concentration of 10 ng/mL (Sigma-Aldrich #GF322) to protect cultures from cell death and support neurite outgrowth. One day after cell plating, FDU was added to inhibit non-neuronal cell proliferation at the final concentration of 6.7 µg/mL. The cultures were imaged immediately or maintained healthy for up to 4-6 weeks.

All cell culture solutions were sterile-filtered (with the pore size of 0.22 µm) before use. The cultures were incubated in a Thermo Scientific Forma Series II water-jacketed incubator at 5% CO_2_ and 37°C. Further details about the preparation of the used reagents are available in Staal et al. (2007).

### 2.2. Neuronal monolayers of the mouse midbrain

These cultures consisted only of a neuronal layer, with no glial layer. The midbrain tissue was dissected from mouse pups as described above. After dissociation, cells were washed, gently triturated with cNSM, and counted. We found that in this step cNSM could be replaced with mNSM (described below) with 1% heat-inactivated fetal bovine serum. Cells were diluted to a density of 1,000,000-1,500,000 cells/mL and plated at around 50,000-100,000 cells per culture dish. Lighter trituration resulted in denser but healthy cultures. The neuron-specific medium (mNSM) consisted of 95% Gibco Neurobasal Plus Medium (ThermoFisher #A3582901), 2% Gibco B-27 Plus Supplement (ThermoFisher #17504044), 2% GlutaMAX (ThermoFisher # 35050061), and 0.5% penicillin-streptomycin (all concentrations are final). The addition of GDNF and FDU, as well as the other procedures, were the same as in the co-cultures. The cultures were imaged immediately or maintained healthy for up to 4-6 weeks.

### 2.3. Immunocytochemistry

Cultures were fixed by aspirating the culture medium and immediately adding phosphate-buffered 4% paraformaldehyde (PFA) for 10 minutes. They were rinsed in 0.1 M phosphate-buffered saline (PBS) and either processed immediately or stored for a few days at 4°C. All immunocytochemical procedures were done at room temperature on a shaker. Cultures were rinsed in PBS, blocked for 15 minutes in 2% normal donkey serum (NDS) in PBS, incubated in goat anti-5-HT IgG (1:1000; ImmunoStar # 20079) and rabbit anti-MAP2 IgG (1:1000; Abcam #32454) with 2% NDS and 0.3% Triton X-100 (TX) in PBS for 1 hour, rinsed 3 times in PBS (5 minutes each), incubated in Cy3-conjugated donkey anti-goat IgG (1:500; ImmunoResearch #705-165-147) and AlexaFluor 488-conjugated donkey anti-rabbit IgG (1:1000; ThermoFisher #A-21206) with 2% NDS in PBS for 30 minutes, and rinsed 3 times (5 minutes each) with PBS. After a quick (5-10 seconds) rinse in water (to remove salts), the coverslip with the cells was carefully detached from the bottom of the plate and mounted on a glass slide with ProLong Gold Antifade Mountant containing DAPI (ThermoFisher #P36941). The cells were imaged at least 24 hours after mounting to allow curing to the optimal refractive index.

### 2.4. Epifluorescence and confocal microscopy

Epifluorescence imaging was performed on the AxioVision Z1 system in three channels (Cy3, GFP, and DAPI), using a 10× objective (NA 0.45) and a 40×oil objective (NA 1.30). Confocal imaging was performed in three channels (Cy3, AlexaFluor 488, DAPI) on the Leica SP8 resonant scanning confocal system, primarily using a 63×oil objective (NA 1.40) with the xy-resolution of 59 nm/pixel and the z-resolution of 300 nm/optical section. Typical z-stacks consisted of 30-100 optical sections. The figures show maximum-intensity projections.

### 2.5. Holotomography (refractive index-based 3D-live imaging)

Live cultures were removed from the incubator and immediately imaged with the 3D-Cell Explorer CX-F (firmware 1.5.70; Nanolive SA, Switzerland) equipped with a 60× objective and a CMOS camera with 1024×1024 pixels. The laser used for tomography was 520 nm at an output power of 0.1 mW. The imaging conditions matched the incubation conditions (5% CO_2_ at 37°C). The z-stacks contained around 95 sections and were imaged around every 7 seconds. The 4D-recordings were reviewed and analyzed in STEVE 1.6 (the native software of the system). Time-lapse z-projections were generated in ImageJ with a Nanolive macro.

### 2.6. Immunohistochemistry and super-resolution microscopy (STED)

A timed-pregnant C57BL/6 dam (Charles River) was euthanized with CO_2_ at embryonic day 17 (E17). The embryos were removed from the uterus, immediately decapitated, and their brains were dissected and immersion-fixed in 4% PFA overnight at 4°C. They were cryoprotected in phosphate-buffered 30% sucrose for 2 days and embedded in 20% gelatin (type A; FisherScientific #G8-500). The gelatin block was trimmed around the brain, immersed in formalin with 20% sucrose for 3 hours, rinsed in PBS, and sectioned on a freezing microtome at 40 µm thickness. Selected sections were rinsed in PBS, blocked in 2% normal goat serum (NGS) in PBS for 30 minutes, and incubated in rabbit anti-5-HT IgG (1:500; ImmunoStar #20080) with 2% NDS and 0.3% TX in PBS for 2 days on a shaker at 4°C. They were rinsed 3 times in PBS (10 minutes each), incubated in STAR-RED-conjugated goat anti-rabbit IgG (1:200, Abberior #STRED-1002) for 90 minutes, rinsed 3 times in PBS (10 minutes each), mounted on coverslips (to minimize the objective-section distance and improve imaging depth), and allowed to air-dry. The coverslips were mounted on glass slides with ProLong Gold Antifade Mountant (without DAPI; ThermoFisher #P36930). The sections were imaged at least 24 hours after mounting to allow curing to the optimal refractive index. They were imaged on the Abberior STED (Stimulated Emission Depletion) microscope using a 60×oil objective (NA 1.4), the excitation line of 640 nm, and the depletion line of 775 nm. The voxel dimensions were 30×30×100 nm^3^. Double-label immunohistochemistry for 5-HT and MAP2 (with AlexaFluor 594-conjugated donkey anti-goat IgG and AlexaFluor 647-conjugated donkey anti-rabbit IgG) was also attempted but yielded virtually no improvement over regular confocal imaging, likely due to suboptimal properties of the AlexaFluor dyes in the used STED configuration.

## 3. RESULTS

Midbrain serotonergic neurons grown in primary cultures were strongly immunoreactive for 5-HT and had normal morphology (Fig. 1). Their somata were round (typically, around 20 µm in diameter) or fusiform (extending up to 50 µm in length) and immunoreactive for MAP2. They typically had long, 5-HT-immunoreactive axons distinguished from other neurites (*e*.*g*., dendrites) by the characteristic varicosity-like profiles and the absence of MAP2-immunoreactivity. Serotonergic neurons appeared normal at various plating densities. In particular, many axons were observed extending from raphe regions that have not been fully dissociated (Fig. 1A), as well as from sparsely distributed single neurons (Fig. 1B-C).

**Figure 1.**
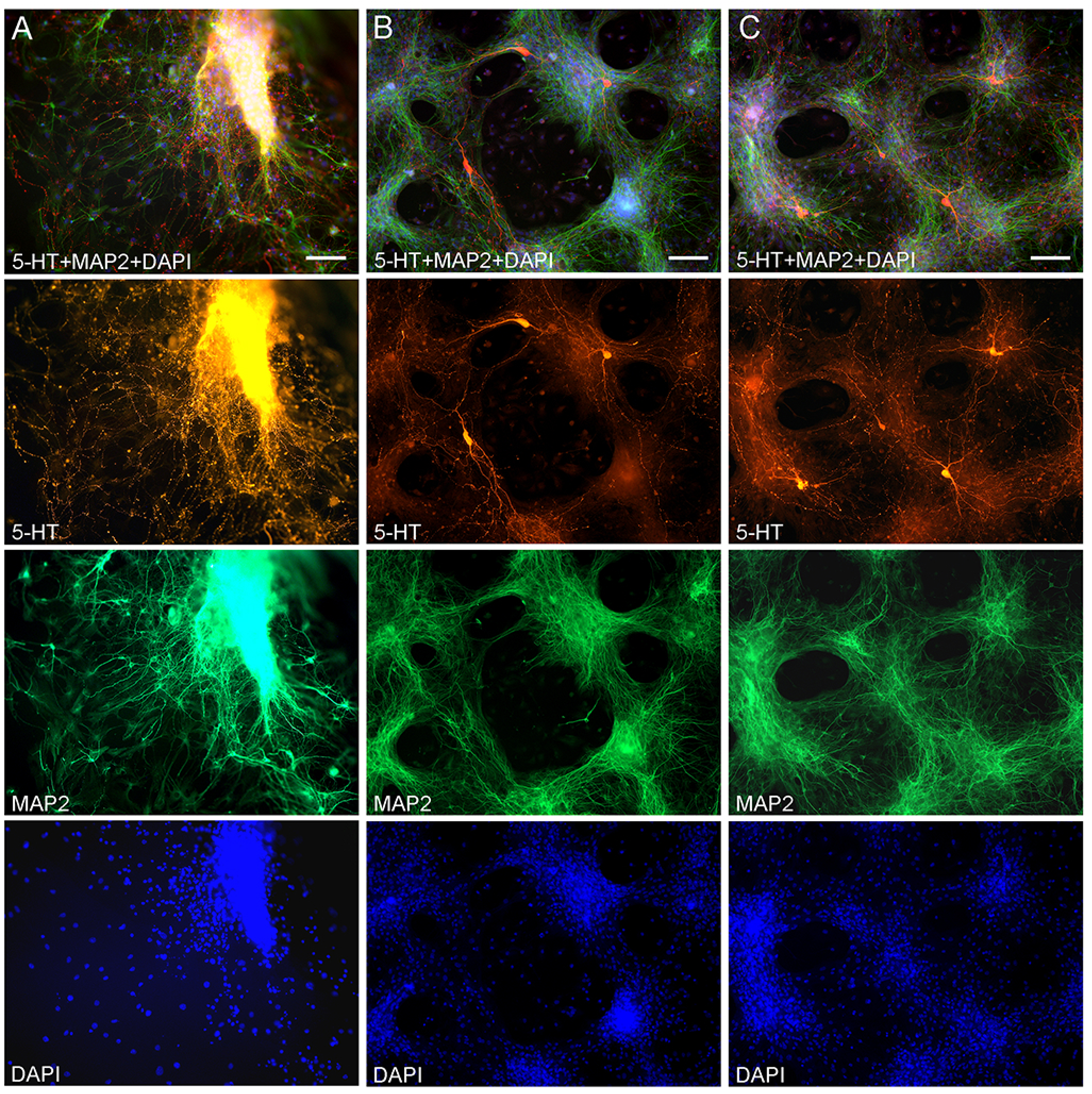
Primary midbrain cultures (all monolayers at DIV 6), visualized with immunocytochemistry for 5-HT (red) and MAP2 (green) and imaged with epifluorescence microscopy. Cell nuclei are stained blue (DAPI). (A) A tissue piece from the midbrain raphe region, the cells of which have not been fully dissociated. Long serotonergic axons with varicosities emerge from the tissue, suggesting that the culture protocol can also be used in organotypic preparations. (B and C) Typical dissociated serotonergic neurons, with morphological features (round or fusiform somata) and neurites virtually indistinguishable from those in intact neural tissue. Scale bar = 100 µm.

High-resolution confocal imaging revealed that many serotonergic axons were in persistent contact with MAP2-positive neurites (*e*.*g*., putative dendrites) of non-serotonergic neurons (Fig. 2). Some serotonergic axons appeared to wind around these neurites (Fig. 2A) and some simply slid along them in the same imaging plane (Fig. 2B). These neurite contacts appeared to be functionally important for axon extension, as was evidenced by instances in which one axon branch remained on the neurite but the other lost its contact (Fig. 2C). In some cases, the detached branch increased its caliber dramatically (at least two-fold), likely due to its multiple active (growth cone-like) zones in the terminal segment. The appearance of these branches in fixed preparations suggested they were attempting to find the next attachment point (Fig. 2C). Serotonergic axons were also found sliding along other serotonergic axons, with no apparent repulsion, but these instances were considerably less frequent (Fig. 2E).

**Figure 2.**
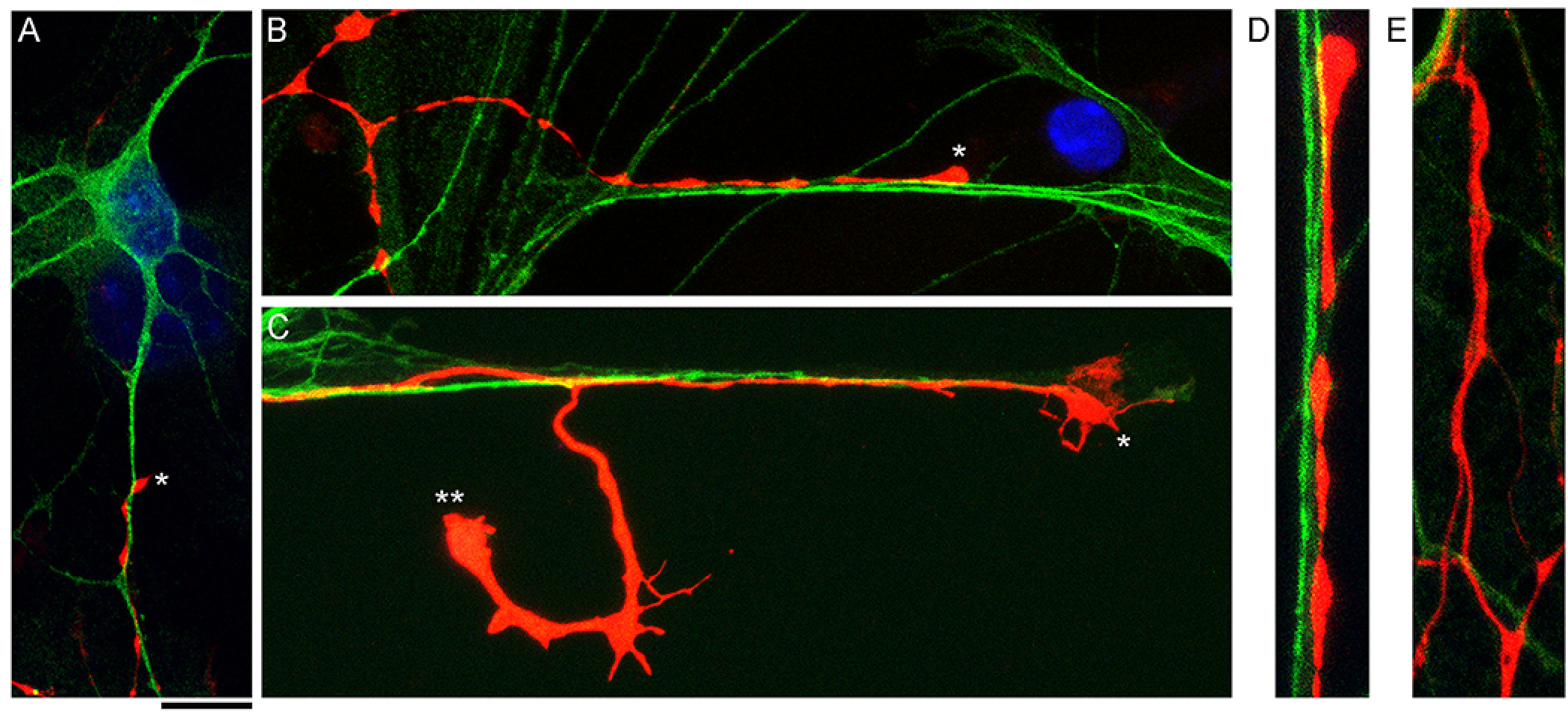
Primary midbrain cultures, visualized with immunocytochemistry for 5-HT (red) and MAP2 (green) and imaged with high-resolution confocal microscopy. Cell nuclei are stained blue (DAPI). (A) A serotonergic axon (5-HT+/MAP2-, asterisk) that advances along a dendrite of a non-serotonergic neuron (5-HT-/MAP2+). (B) Another serotonergic axon (5-HT+/MAP2-, asterisk) that advances along a 5-HT-/MAP2+ neurite. (C) A serotonergic axon (5-HT+/MAP-, asterisk) that advances to the end of a 5-HT/MAP2+ neurite and also produces a branch that has lost contact with the neurite (double asterisk). Note the much larger caliber of the branch, as well as multiple growth cone-like zones, perhaps in search of the next attachment point. (D) A typical contact between a serotonergic axon (5-HT+/MAP2-) and a 5-HT-/MAP2+ neurite (an enlarged part of B). (E) Contacts between two serotonergic axons (5-HT+/MAP2-) are less frequent but also occur, with no apparent repulsion between the axons. A, B, D, and E: monolayers at DIV 4; C: neuron-glia co-culture at DIV 3. Scale bar (shared by all panels) = 10 µm in A, B and C; 5 µm in D and E.

In order to investigate the nature of contacts between serotonergic axons and MAP-positive neurites, we examined axons that were advancing along a neurite but were not in close apposition to it (Fig. 3A). It revealed discrete adhesion structures, composed of a strongly 5-HT-postive “foot,” directly in contact with the neurite, and an extremely thin (nanoscale) membrane tether anchoring it to the main axon (Fig. 3B-C). These structures were located close to the growth cone but outside its active zone. Putative adhesion sites were also detected on neurites where the axon was no longer present. In some instances, these sites were strongly elongated (Fig. 3D), suggesting that the contacting axon membrane was flattened, perhaps to accommodate the width of the neurite.

**Figure 3.**
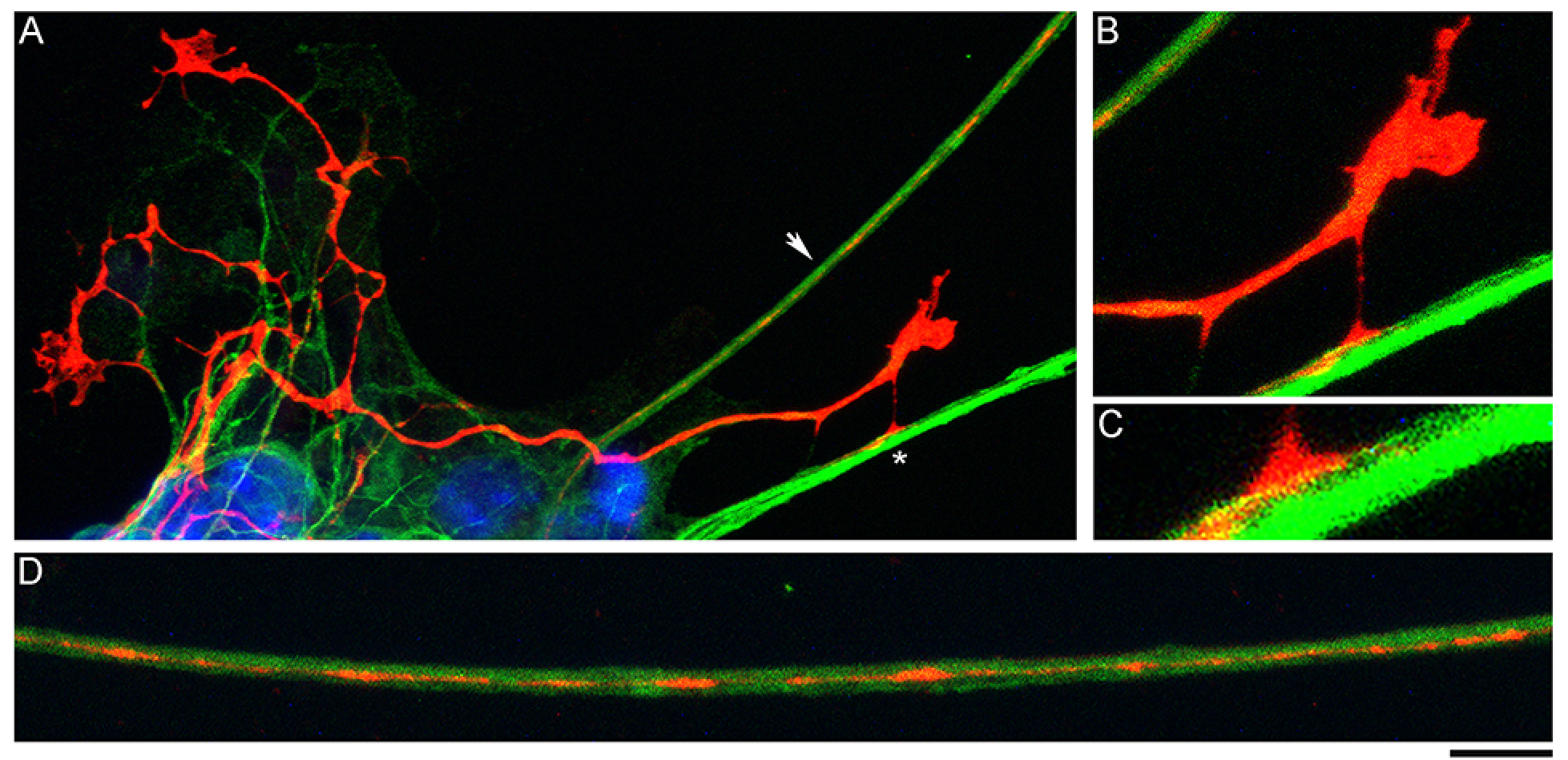
A primary midbrain culture (neuron-glia co-culture at DIV 3), visualized with immunocytochemistry for 5-HT (red) and MAP2 (green) and imaged with high-resolution confocal microscopy. Cell nuclei are stained blue (DAPI). (A) A serotonergic axon (5-HT+/MAP2) that advances along a 5-HT-/MAP2+ neurite and reveals its adhesion sites (one site is marked with an asterisk). (B) An enlarged view of the growth cone region in A. (C) A further enlarged view of the adhesion site marked with an asterisk in A. (D) A relatively regularly spaced putative adhesion sites (5-HT+/MAP2-) on a 5-HT-/MAP2+ neurite (an enlarged part of A, arrow). Scale bar (shared by all panels) = 10 µm in A; 5 µm in B and D; 2 µm in C.

The presence of 5-HT-positive adhesion sites raises the question of whether they can be mistaken for varicosities in brain tissue. Image analysis suggests subtle transitions between actual axon varicosities (connected with membrane) and residual adhesion sites (with no membrane continuity), which may be difficult to distinguish in fixed tissue visualized with immunohistochemistry (Fig. 4). This situation is further complicated by the observation that serotonergic axons themselves can have segments with no detectable 5-HT-immunoreactivity, despite their large caliber and strong immunoreactivity in other segments (Fig. 4A). This phenomenon may reflect the typical formation of serotonergic varicosities, by either the accumulation of 5-HT in specific segments or the production of extremely thin inter-varicose segments that fall below the limit of optical resolution. However, it is also possible that distal axon segments can become detached from the main axon, perhaps due to the highly dynamic active zones, extending beyond the growth cone (*e*.*g*., Fig. 2C). Since these zones can be accompanied by rapid fluctuations of the axon diameter, distal segments may become disconnected in stochastic events. Such terminal “shedding” would not be detrimental to neural tissue because serotonergic axons exhibit robust growth and naturally regenerate, even in the adult mammalian brain (Jin et al., 2016).

**Figure 4.**
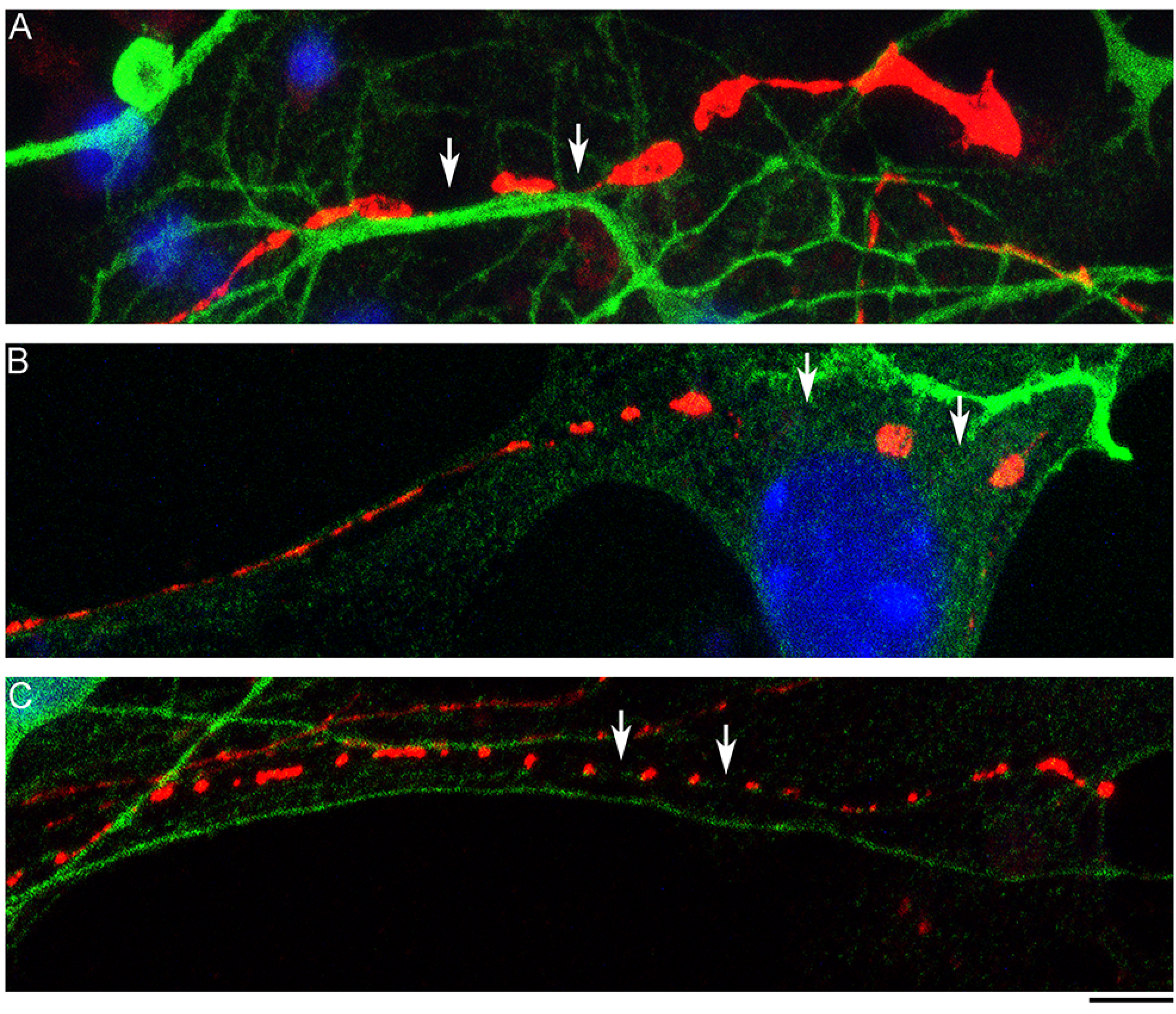
Primary midbrain cultures (all monolayers at DIV 4), visualized with immunocytochemistry for 5-HT (red) and MAP2 (green) and imaged with high-resolution confocal microscopy. (A) The growth-cone region of a serotonergic axon (5-HT+/MAP2-) that shows wide gaps in 5-HT-immunoreactivity (arrows). These gaps may be due to nanoscale-caliber bridges between intensely labeled segments or actual interruptions in fiber continuity. The presence of normal-caliber segments virtually devoid of 5-HT also cannot be ruled out. (B) A serotonergic axon or a series of its adhesion sites (5-HT+/MAP2-) that transitions from a thin, nearly continuous trace to a much thicker trace with large gaps (arrows) between circular 5-HT+ regions. (C) A serotonergic axon or a series of its adhesion sites (5-HT+/MAP2-) that shows circular 5-HT+ regions (around 1 µm in diameter) spaced at around 4 µm (arrows). Scale bar (shared by all panels) = 5 µm.

In order to understand the dynamics of these processes, midbrain cultures were examined with time-lapse 3D-holotomography (label-free, refractive index-based imaging). Highly dynamic, discrete contact events between growth-cone protrusions and neurites were recorded (Fig. 5). Some of these contacts may eventually become adhesion sites (Fig. 6). As the axon continues to advance along the surface, its spatial position retains a considerable degree of flexibility. Specifically, it can transition from being closely adhered to the surface (*e*.*g*., Fig. 2B) to being anchored only by thin membrane tethers (*e*.*g*., Fig. 3B). Live imaging suggests that these tethers can extend by around 10 µm (the diameter of a small neuron) in around 10 minutes, while maintaining the attachment site firmly fixed (Fig. 6). Such physical flexibility is important to accommodate unavoidable lateral shifts of the axon, as its leading end clambers from neurite to neurite (Fig. 7). Perhaps in response to tension forces, axon segments can be come flat and assume a cork screw-like configuration, even far away from the growth cone and naturally flat lamellipodia (Fig. 6, inset).

**Figure 5.**
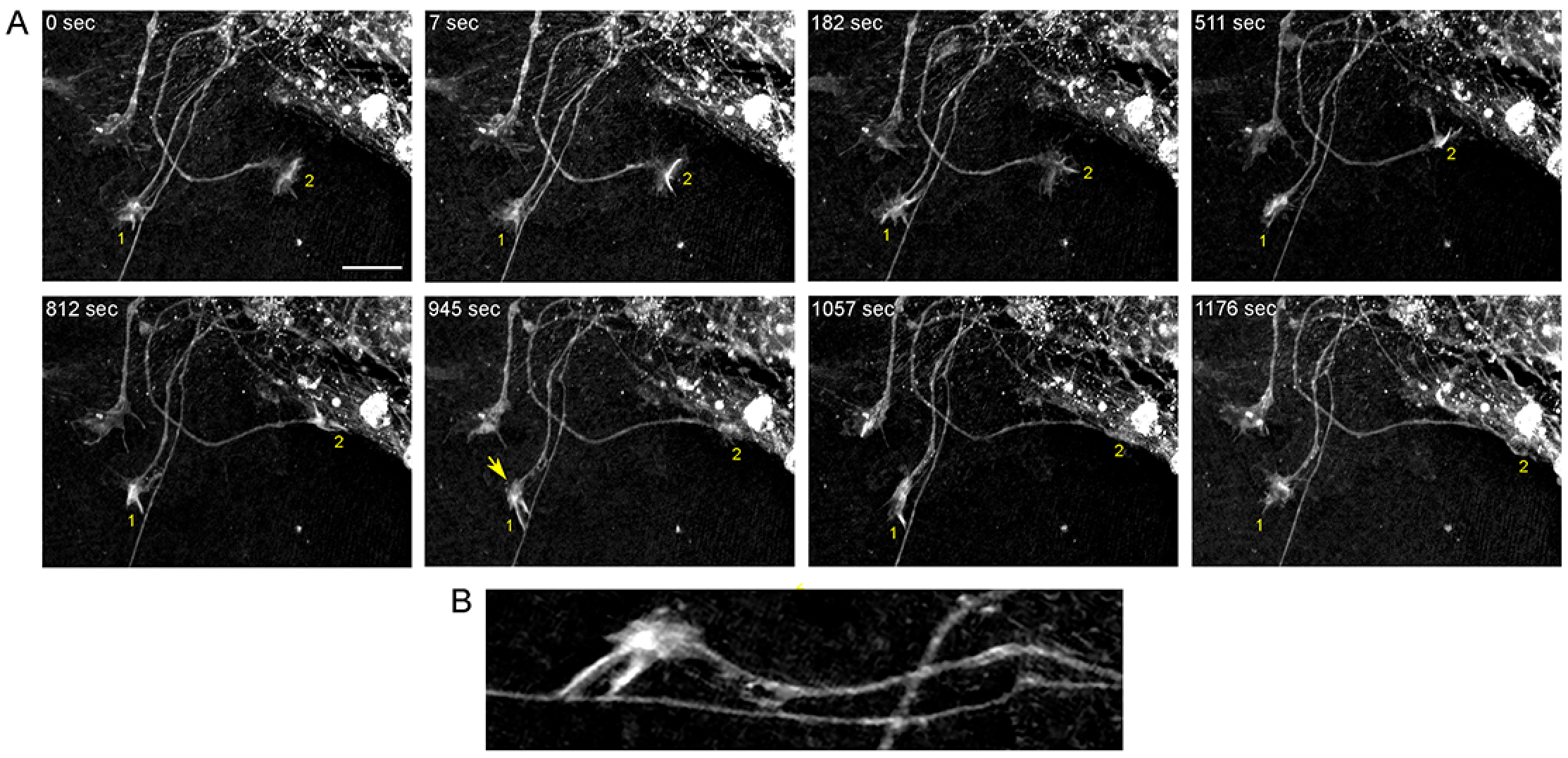
The dynamics of two growth cones (GC; labeled 1 and 2) in a primary midbrain culture (monolayer at DIV 5), visualized with time-lapse holotomography. (A) GC 1 attempts to move along a neurite by producing protrusions that come in contact with the neurite at well-defined points (arrow, enlarged in B). The distance between adjacent points is similar to that between the discrete 5-HT+ regions in Figure 4. GC 2 detects a substrate (around *t* = 511 sec) and rapidly advances along its edge. To emphasize key transitions, time points are not evenly spaced. Scale bar = 10 µm.

**Figure 6.**
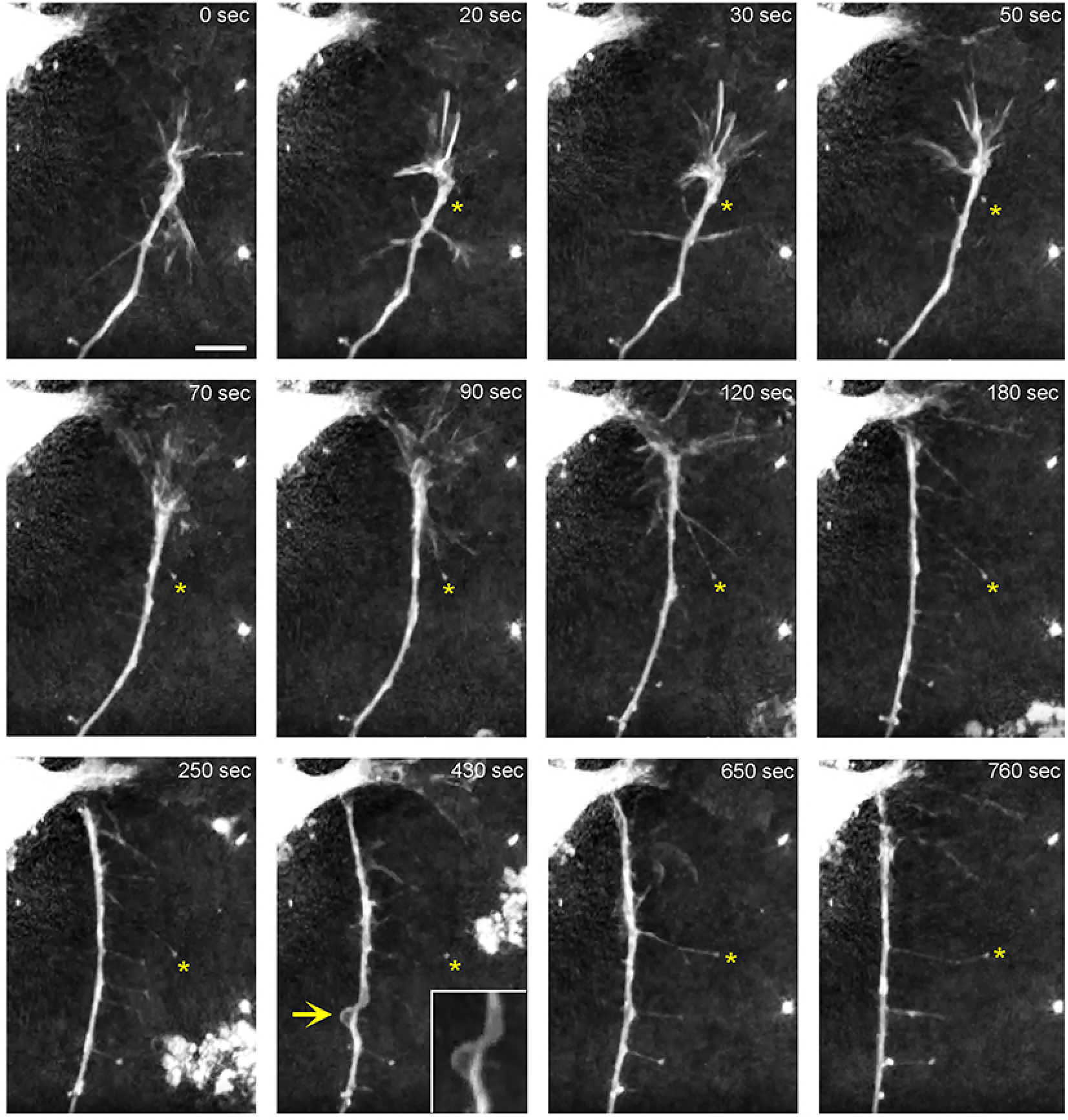
The dynamics of a growth-cone region in a primary midbrain culture (monolayer at DIV 1), visualized with time-lapse holotomography. As the growth cone finds its next attachment target and pulls the axon to the left, its adhesion sites are revealed (one such site is marked with an asterisk). The spacing between the adhesion sites is around 2-6 µm, and the nanoscale tethers connecting them to the main axon can be stretched to as long as 10 µm (to accommodate axon shifts). This spatial configuration closely resembles that shown in Figure 3. The arrow points to an axon segment that appears flat and twisted in a corkscrew-like fashion. To emphasize key transitions, time points are not evenly spaced. Scale bar = 5 µm.

**Figure 7.**
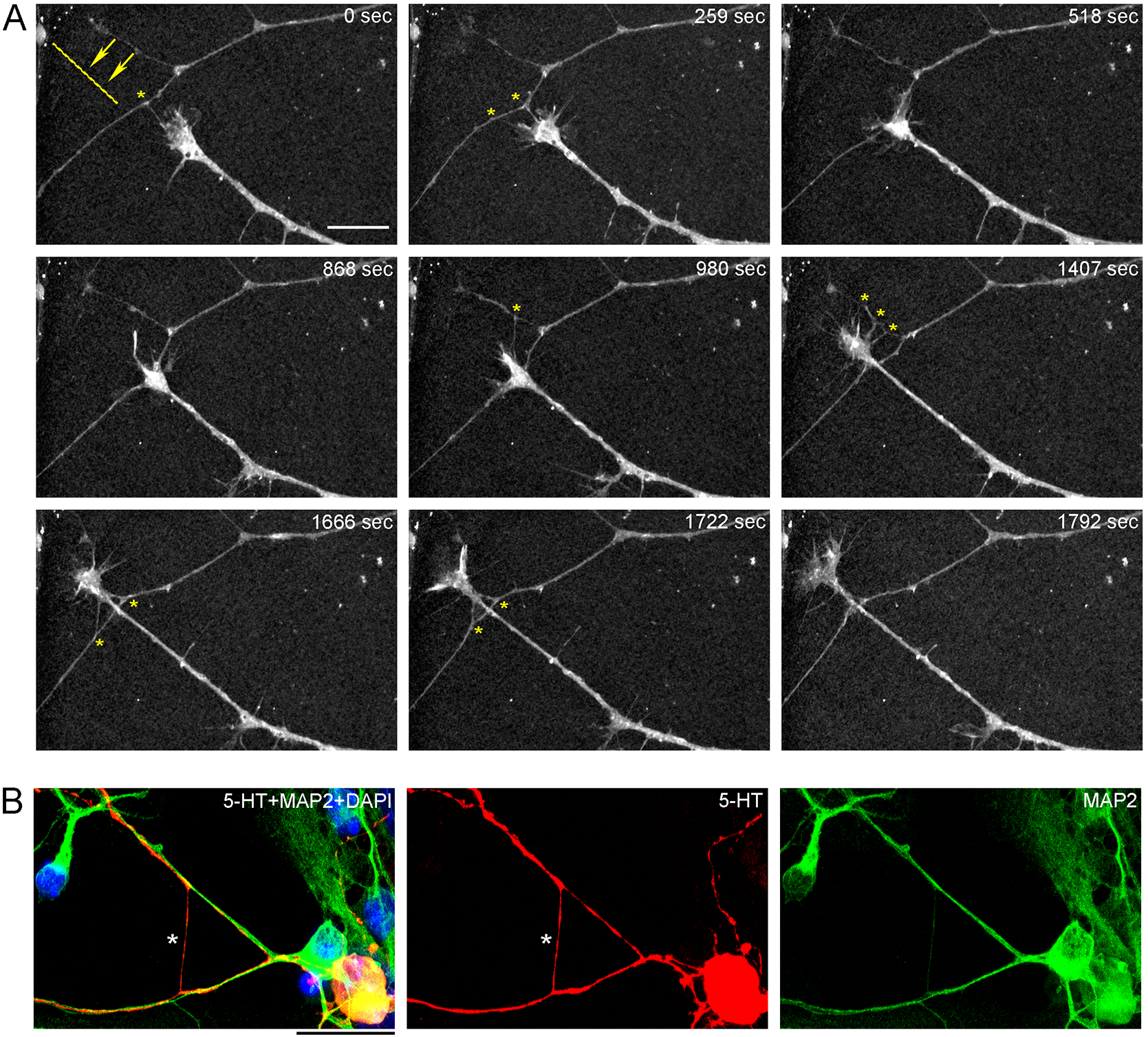
(A) The dynamics of a growth-cone region in a primary midbrain culture (monolayer at DIV 5), visualized with time-lapse holotomography. A transition from one neurite branch to another is shown (some key adhesion points are marked with asterisks). The arrows with the dashed line indicate the spatial shift of the top branch, as the growing axon generates sufficient force to line it up with its current growth axis. To emphasize key transitions, time points are not evenly spaced. Scale bar = 10 µm. (B) A similar transition (asterisk) in a primary midbrain culture (monolayer at DIV 2), visualized with immunocytochemistry for 5-HT (red) and MAP2 (green) and imaged with high-resolution confocal microscopy. Cell nuclei are stained blue (DAPI). Scale bar = 20 µm.

The environment in the culture dish does not accurately reflect the environment in the brain. The differences include tissue dimensionality (2D vs. 3D), cell packing, viscoelasticity, and many other factors. In order to verify some of our findings, super-resolution (STED) microscopy was used to examine single serotonergic axons in the mouse brain at embryonic day 17 (Fig. 8). This developmental age is convenient because at this time the serotonergic neurons are already fully mature but their axons only begin to spread in the telencephalon (Lidov and Molliver, 1982; Janušonis et al., 2004). Due to this sparse distribution, single axons and their growth cones can be easily captured in natural brain tissue. In the embryonic telencephalon, growth cone protrusions that closely resemble those in culture (*e*.*g*., Fig. 3) were detected. In particular, some of them appeared to have a “foot” and a tether (Fig. 8, inset). In addition, unambiguously flattened membrane segments were detected (with the ratio of approximately 5:1), with an apparent cork-screw rotation (Fig. 9). This suggests that serotonergic axons can be ribbon-like (or human palm-like, to better approximate the ratio) as they travel through neural tissue. Since these profiles were also observed in culture (Figs. 3D, 4B, 6), they may not be induced by dense tissue packing.

**Figure 8.**
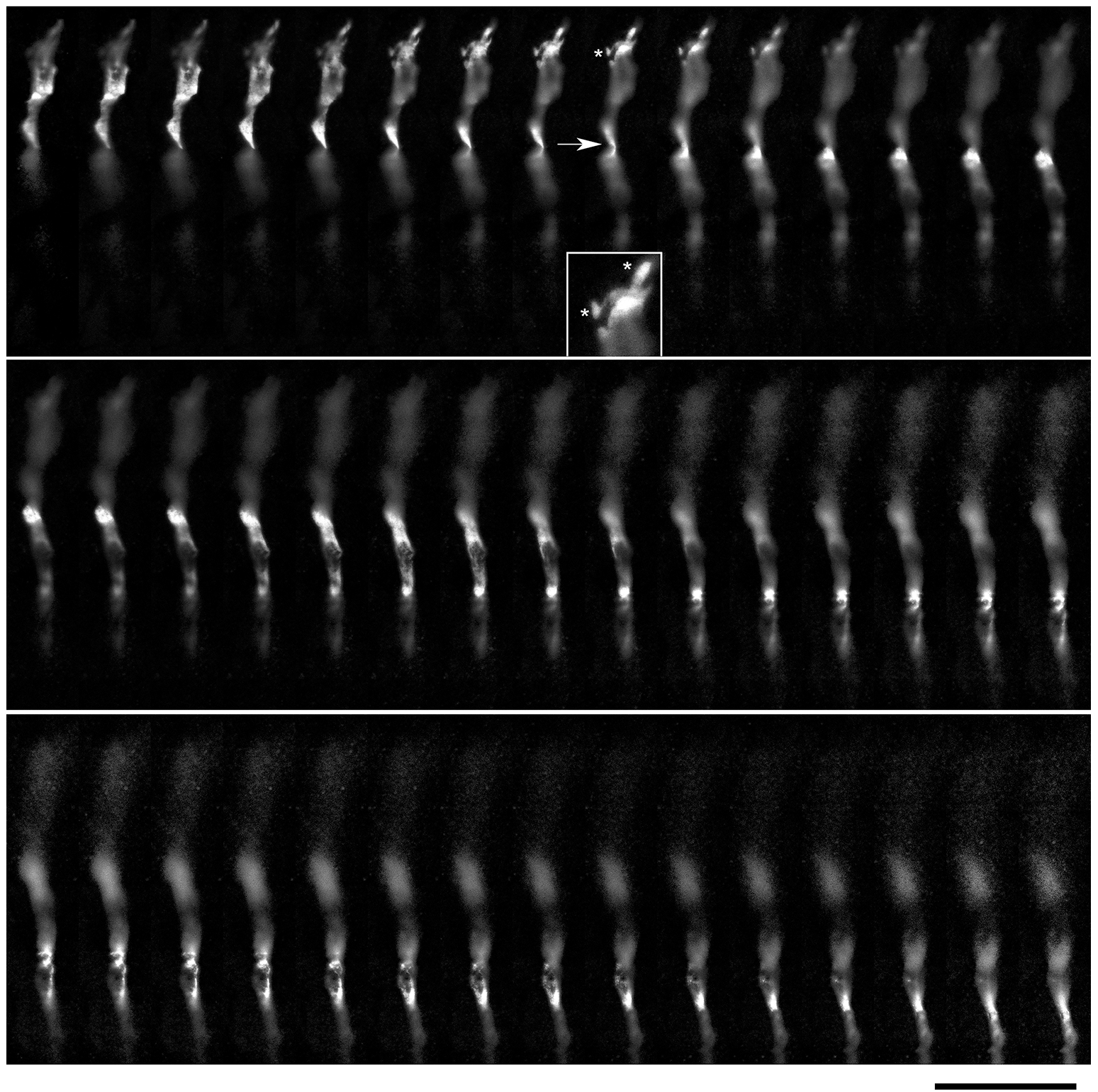
A super-resolution microscopy (STED) z-series of a single serotonergic (5-HT+) axon in the sectioned cortical plate of a mouse embryo at E17. Note the growth cone protrusions similar to those in culture (e.g., Figs. 3B and 10D; asterisks, inset) and a potentially flat membrane region (arrow, further analyzed in Fig. 9). The sequential optical sections are evenly separated by 100 µm. Scale bar = 20 µm.

**Figure 9.**
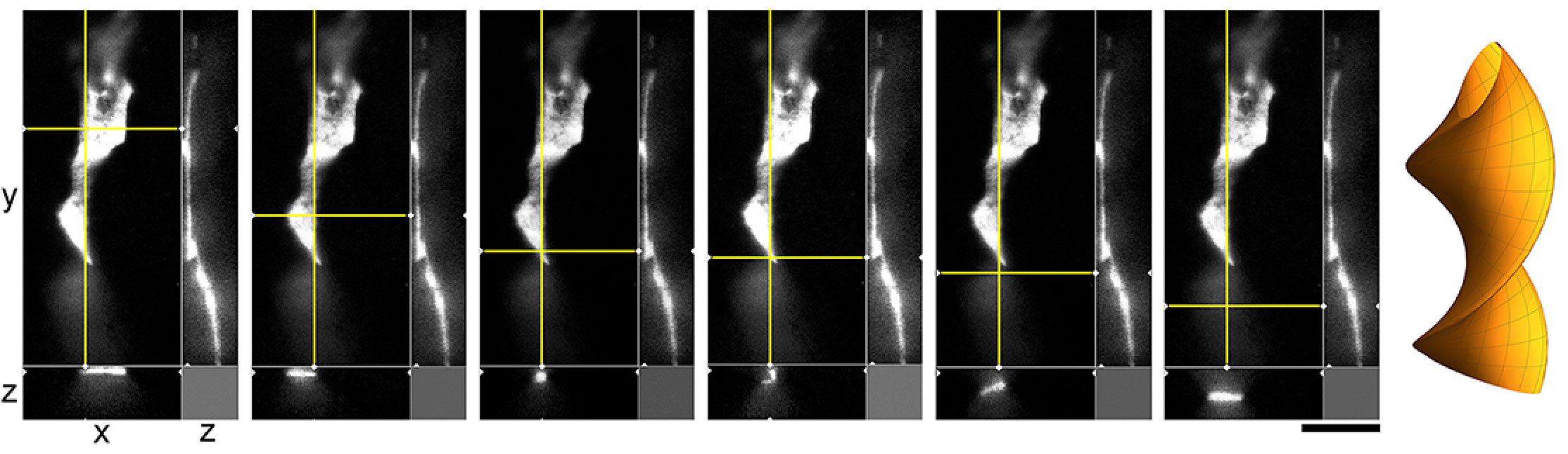
A super-resolution microscopy (STED) images of a single serotonergic (5-HT+) axon in the sectioned cortical plate of a mouse embryo at E17. The axon is shown in all three dimensions, with a fixed yz-plane (at a constant x; the vertical yellow line) and a series of xz-planes (with six different y values; the horizontal yellow line). At the shown levels, the axon is flat (ribbon-like) and appears to rotate. This hypothetical cork-screw configuration is shown in the diagram on the right. Scale bar = 5 µm.

Realistic modeling of serotonergic axons requires experimental information about their branching patterns. At a minimum, it should include the frequency of branching, the typical branching angles, and the trajectory information retained by each of the two branches. This information should ideally be described probabilistically (*e*.*g*., the “wait time” between two branching events might be captured by the exponential distribution with a given intensity λ, the branching angles can be described by a directional probability distribution, and the “memory” of the branches can be reflected in the underlying increment covariance structure). Serotonergic axon ramification is often referred to in descriptive density studies, but this process is essentially inferred, with no reliable information at the level of single axons. In brain tissue, these axons can be extremely dense, to the extent that even high-resolution 3D-imaging can be insufficient to distinguish true branching points from axons that cross at sub-micrometer distances (Janušonis et al., 2019). Serotonergic neurons in culture tended to produced branching events that could be detected unambiguously (Fig. 10). Locally, they appeared to be rather stereotypic. The two branches of an axon split at wide angles (typically, 90°-180°), which achieved their immediate separation (Fig. 10). In sparse cultures, both branches can reorient themselves parallel to the original trajectory and thus cannot be treated as independent of their parent trajectory or of each other. Over longer distances, they are likely to completely decorrelate, as they advance through the stochastically distributed attachment surfaces.

**Figure 10.**
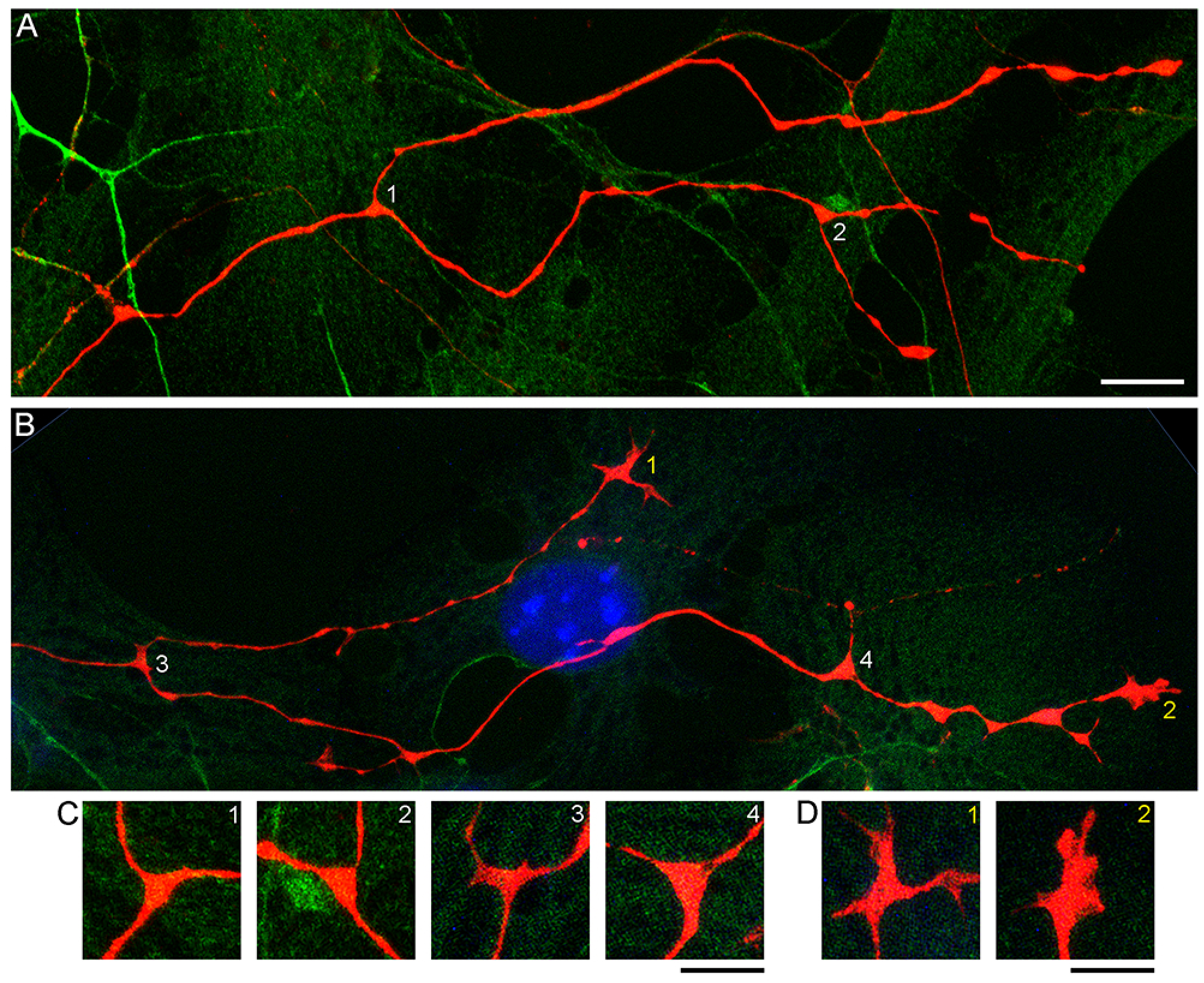
Primary midbrain cultures (monolayers at DIV 4), visualized with immunocytochemistry for 5-HT (red) and MAP2 (green) and imaged with high-resolution confocal microscopy. Cell nuclei are stained blue (DAPI). (A and B) Four branching regions (white 1-4) and two growth cones (yellow 1-2) and are shown. (C) An enlarged view of the branching points. The branching regions have a triangular shape, and the branches diverge at large angles (around 90°-180°). (D) An enlarged view of the growth cones. Note the similarity of the second growth cone to that in a sectioned mouse brain (Fig. 8). Scale bars = 10 µm in A and B, 5 µm in C and D.

## 4. DISCUSSION

This study is the first high-resolution analysis of single serotonergic axons *in vitro*. A key advantage of cell cultures is that long axons can be imaged uninterrupted from the soma to the terminal point, including branching points along the trajectory. In contrast, brain tissue is extremely densely packed, with extracellular distances often well below 100 nm (Hrabetova et al., 2018). This compressed environment may conceal the inherent properties of axons (*e*.*g*., their caliber dynamics), as well as their contacts with other cells. Also, only relatively short axon segments can be visualized uninterrupted in sectioned brain tissue, because their random walk-like trajectories stay within the section volume only for distances comparable to the width of the section (typically, 40-50 µm) (Janušonis et al., 2019). In brain tissue, reliable identification of branching events in serotonergic axons is currently possible only in relatively sparse regions and with sub-micrometer 3D-imaging, due to the fact that serotonergic axons routinely cross at distances near the limit of optical resolution (Janušonis et al., 2019). Therefore, the true extent of branching or ramification in serotonergic axons remains unknown, even though these processes are often used to explain regional density differences.

Cell cultures are limited in that they cannot reproduce the natural environment of cells. In particular, traditional cell cultures and natural tissues differ in their dimensionality (2D vs. 3D) and a number of physical properties that cells respond to (Gjorevski et al., 2022). The burgeoning field of biocompatible hydrogels is already providing a more graded spectrum of experimental platforms between these two extremes (Wisdom and Chaudhuri, 2017; Sullivan et al., 2022). The transcriptomes of neurons in the brain and culture environments are expected to differ; primary cultures of midbrain neurons have been shown to have upregulated expression of genes associated with extracellular matrix and adhesion (Greco et al., 2009). Cultured neurons can also self-organize into functional ensembles (Antonello et al., 2022) that may differ from mesoscale circuits in brain tissue.

Axon extension is a fundamental process that is unlikely to be strongly altered in culture environments, especially if the soma and neurite morphologies do not deviate significantly from those observed in natural neural tissue. Additionally, findings in culture systems can be verified by targeted high-resolution analyses in the brain, whenever feasible.

By taking this approach, we show that serotonergic axons can be ribbon-like (with the width-thickness ratio of around 5:1) and can also rotate along their axis, perhaps producing periodic point-like constrictions (Figs. 8, 9). This morphology is likely induced by tension forces due to axon extension, but it also increases the surface-to-volume ratio and may facilitate 5-HT release. It is less likely to be related to tissue packing because it was also observed in sparse cultures (Fig. 6). Interestingly, these profiles were noted in early studies (see Fig. 5 of (Aitken and Tork, 1988)), where they were interpreted as “sinuous fibers” with “translucent varicosities” (because of their higher light intensity, likely due to the shorter light path in the flat region). It highlights the limited understanding of what serotonergic “varicosities” are, despite their importance in neuroanatomical and functional studies (*e*.*g*., they have been used to classify serotonergic axons into the D- and M-classes (Kosofsky and Molliver, 1987), a system still referred to in some current analyses). Since early descriptions, based on 2D-microscopy, they have been assumed to be dilated segments of axons, but very few studies have performed their high-resolution analyses in the 3D-space. The emerging picture of varicosity-like segments (VLSs) is considerably more complex. First, some VLSs may indeed be dilated, ovoid-shaped segments (Maddaloni et al., 2017). This is supported by our findings that demonstrate that the caliber of the same axon can vary considerably over short distances (*e*.*g*., Fig. 2C). Second, a sequence of VLSs may not indicate significant changes in axon morphology but may reflect a ribbon-like segment that periodically exposes its flat surface versus its edge, effectively creating a “blinking” effect in 2D-imaging. Third, some VLSs may not be continuous axons but rather 5-HT-positive “footprints,” former adhesion sites left by axons. Our study clearly demonstrates this possibility *in vitro*, but additional studies are needed in brain tissue to prove the absence of membrane bridges between adjacent VLSs (*e*.*g*., using multiple axonal markers and super-resolution microscopy). As more experimental information becomes available, VLSs may provide important insights into axonal *dynamics* in *fixed* brain tissue. These observations are generally consistent with recent findings in other systems, where VLSs and other morphological features have been shown to reflect the current, local state of an axon rather than its identity (Liu and Nakamura, 2006; Andersson et al., 2020; Sun et al., 2022).

Our analysis sheds new light on the dispersal of serotonergic axons. It has recently been shown that this process is aided by *Pcdh-αC2* (a protocadherin) that is expressed in serotonergic neurons and can mediate axonal self-avoidance, preventing axon “clumping” (Katori et al., 2009; Chen et al., 2017; Katori et al., 2017). However, our previous modeling has shown that “clumping” is unlikely if axons perform random walks with no interaction (based on the von Mises-Fisher step-wise walk with a high concentration parameter (κ) or fractional Brownian motion with a high Hurst index (*H*)) (Janušonis and Detering, 2019; Janušonis et al., 2020). In this context, branching points can be disruptive because they can create sister trajectories that travel in close proximity, at least initially. *In vitro* results demonstrate that the sister branches of serotonergic axons tend to separate at very large angles, which efficiently prevents clustering. In the brain, each of the branches is likely to immediately encounter different adhesion surfaces, supporting rapid decorrelation.

An important finding in this study is that serotonergic axons prefer to travel adhered to available cellular surfaces, such as dendritic branches of other neurons. Some of the key axonal structures supporting this adhesion might have been observed in early studies but could not be accurately interpreted because of technical limitations (Azmitia and Whitaker-Azmitia, 1987; Aitken and Tork, 1988). In the densely packed neural tissue, dendrites and other cellular surfaces are readily available. This suggests that the strong stochasticity of serotonergic axon trajectories may not be a property of serotonergic fibers themselves but may instead reflect the stochastic geometry of the surrounding neural tissue (Fig. 11). Given a specified stochastic process, computer simulations can predict the resultant axon densities, with no additional biological information (Janušonis and Detering, 2019; Janušonis et al., 2020). This leads to an intriguing conclusion that local serotonergic densities can directly reflect the local microarchitecture of a brain region. In particular, an abnormal local cytoarchitecture alone may generate an altered serotonergic density, with no other causal factors. It might be exemplified by the increased densities of serotonergic axons in some cortical regions of individuals with Autism Spectrum Disorder (Azmitia et al., 2011), perhaps in association with the reported denser cell packing in cortical minicolumns (Casanova et al., 2006).

**Figure 11.**
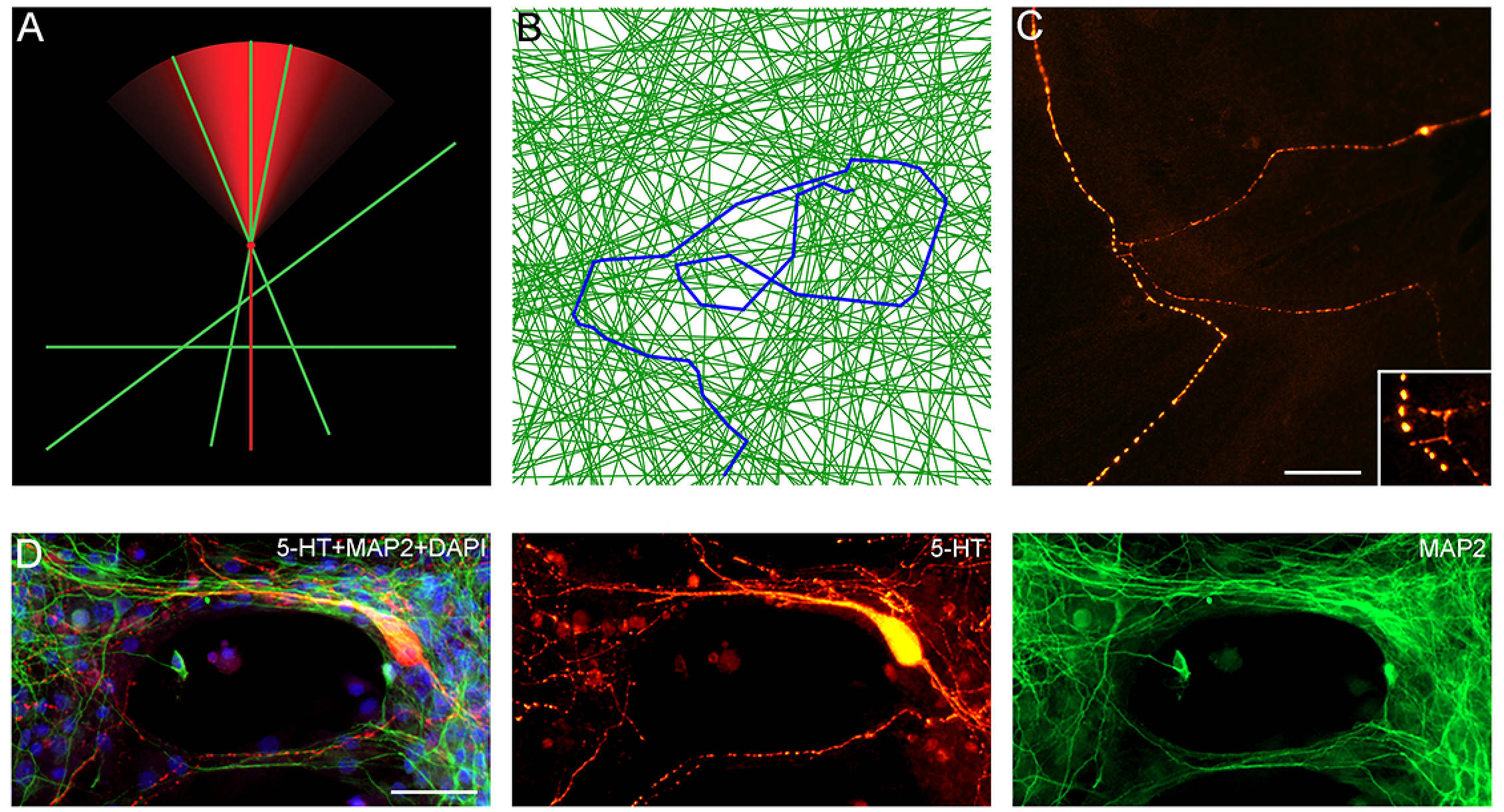
(A) A diagram showing the hypothetical choices available to an advancing serotonergic axon (red) that can choose any of the available neurites (green), provided they fall within its sector of possibilities. The cone is modeled with the von Mises distribution (here, the mean direction **µ** is aligned with the current direction of the fiber and the concentration parameter κ = 10. (B) A simulated walk of a fiber (blue) that can only advance only along available neurites (green). In the simulation, 200 randomly-oriented lines were used, and the fiber advanced through 350 intersections. At each intersection, it could move in the current direction (a 0°-turn), turn “left” or “right” (at the available angles), or turn backward (a 180°-turn). The probabilities of the four events were calculated using the von-Mises distribution with **µ** = (0, 0) and κ = 5, and one event was drawn. The simulation was performed in Wolfram Mathematica 13.0. (C) A comparable configuration of 5-HT+ axons in a glia-neuron coculture at DIV 15, visualized with epifluorescence microscopy. The inset shows potential contact points. Scale bar = 20 µm. (D) Serotonergic axons (5-HT+/MAP2-, red) traveling along bridges of MAP2+ (green) neurites, visualized with epifluorescence microscopy. Scale bar = 50 µm.

The rapid expansion of the currently available toolbox, including approaches developed in our research program (holotomography of primary brainstem cultures, advanced stochastic modeling, supercomputing simulations), promises to produce a radically new view of the serotonergic system, both at the structural and functional levels. The recent development of pluripotent stem cell-derived human serotonergic neurons (Lu et al., 2016; Cao et al., 2017; Vadodaria et al., 2018), combined with these methods, will also stimulate novel theory-guided approaches to brain restoration after injury.

## 6. Author Contributions

MH developed all midbrain cell cultures, directed all technical improvements, and contributed to all theoretical discussions (from data interpretation to modeling applications). AMLV and JES made major contributions in assisting MH in the initial development of the system. SJ initiated and supervised the project within the novel conceptual framework of stochastic axon systems.

## 8. Acknowledgements

We thank Dr. David Sulzer and Ellen Kanter (Columbia University) for their extensive technical advice in setting up a midbrain *in vitro* system at UCSB. We acknowledge the contributions of Hao Li, Justin Chiu, Samantha Siu, and Allison Vargas (members of the Janušonis laboratory). We also thank Dr. Benjamin Lopez (UCSB Neuroscience Research Institute) and Dr. Jennifer Smith (UCSB California NanoSystems Institute) for their technical support and the facilitation of the imaging logistics.

## 9. Data Availability Statement

All computer code and original images are available on request from the corresponding author.

## REFERENCES

Adori, C., Ando, R.D., Szekeres, M., Gutknecht, L., Kovacs, G.G., Hunyady, L., et al. (2011). Recovery and aging of serotonergic fibers after single and intermittent MDMA treatment in Dark Agouti rat. J Comp Neurol 519(12), 2353–2378. doi: 10.1002/cne.22631.

Aitken, A.R., and Tork, I. (1988). Early development of serotonin-containing neurons and pathways as seen in wholemount preparations of the fetal rat brain. J Comp Neurol 274(1), 32–47. doi: 10.1002/cne.902740105.

Andersson, M., Kjer, H.M., Rafael-Patino, J., Pacureanu, A., Pakkenberg, B., Thiran, J.P., et al. (2020). Axon morphology is modulated by the local environment and impacts the noninvasive investigation of its structure-function relationship. Proc Natl Acad Sci USA 117(52), 33649–33659. doi: 10.1073/pnas.2012533117.

Antonello, P.C., Varley, T.F., Beggs, J., Porcionatto, M., Sporns, O., and Faber, J. (2022). Self-organization of in vitro neuronal assemblies drives to complex network topology. Elife 11. doi: 10.7554/eLife.74921.

Azmitia, E.C., Singh, J.S., and Whitaker-Azmitia, P.M. (2011). Increased serotonin axons (immunoreactive to 5-HT transporter) in postmortem brains from young autism donors. Neuropharmacology 60(7-8), 1347–1354. doi: 10.1016/j.neuropharm.2011.02.002.

Azmitia, E.C., and Whitaker-Azmitia, P.M. (1987). Target cell stimulation of dissociated serotonergic neurons in culture. Neuroscience 20(1), 47–63. doi: 10.1016/0306-4522(87)90005-4.

Butler, A.B., and Hodos, W. (2005). Comparative Vertebrate Neuroanatomy (2nd edition). Wiley-Interscience.

Campanelli, F., Marino, G., Barsotti, N., Natale, G., Calabrese, V., Cardinale, A., et al. (2021). Serotonin drives striatal synaptic plasticity in a sex-related manner. Neurobiol Dis 158, 105448. doi: 10.1016/j.nbd.2021.105448.

Cao, L., Hu, R., Xu, T., Zhang, Z.N., Li, W., and Lu, J. (2017). Characterization of Induced Pluripotent Stem Cell-derived Human Serotonergic Neurons. Front Cell Neurosci 11, 131. doi: 10.3389/fncel.2017.00131.

Carrera, I., Molist, P., Anadon, R., and Rodriguez-Moldes, I. (2008). Development of the serotoninergic system in the central nervous system of a shark, the lesser spotted dogfish Scyliorhinus canicula. J Comp Neurol 511(6), 804–831. doi: 10.1002/cne.21857.

Casanova, M.F., van Kooten, I.A., Switala, A.E., van Engeland, H., Heinsen, H., Steinbusch, H.W., et al. (2006). Minicolumnar abnormalities in autism. Acta Neuropathol 112(3), 287–303. doi: 10.1007/s00401-006-0085-5.

Casper, D., Mytilineou, C., and Blum, M. (1991). EGF enhances the survival of dopamine neurons in rat embryonic mesencephalon primary cell culture. J Neurosci Res 30(2), 372–381. doi: 10.1002/jnr.490300213.

Chen, W.V., Nwakeze, C.L., Denny, C.A., O’Keeffe, S., Rieger, M.A., Mountoufaris, G., et al. (2017). Pcdhalphac2 is required for axonal tiling and assembly of serotonergic circuitries in mice. Science 356(6336), 406–411. doi: 10.1126/science.aal3231.

Cooke, P., Janowitz, H., and Dougherty, S.E. (2022). Neuronal Redevelopment and the Regeneration of Neuromodulatory Axons in the Adult Mammalian Central Nervous System. Front Cell Neurosci 16, 872501. doi: 10.3389/fncel.2022.872501.

Daws, R.E., Timmermann, C., Giribaldi, B., Sexton, J.D., Wall, M.B., Erritzoe, D., et al. (2022). Increased global integration in the brain after psilocybin therapy for depression. Nat Med 28(4), 844–851. doi: 10.1038/s41591-022-01744-z.

di Porzio, U., Daguet, M.C., Glowinski, J., and Prochiantz, A. (1980). Effect of striatal cells on in vitro maturation of mesencephalic dopaminergic neurones grown in serum-free conditions. Nature 288(5789), 370–373. doi: 10.1038/288370a0.

Donovan, L.J., Spencer, W.C., Kitt, M.M., Eastman, B.A., Lobur, K.J., Jiao, K., et al. (2019). Lmx1b is required at multiple stages to build expansive serotonergic axon architectures. Elife 8, e48788. doi: 10.7554/eLife.48788.

Ebbesson, L.O., Holmqvist, B., Ostholm, T., and Ekström, P. (1992). Transient serotonin-immunoreactive neurons coincide with a critical period of neural development in coho salmon (Oncorhynchus kisutch). Cell Tissue Res 268(2), 389–392. doi: 10.1007/bf00318807.

Fabbiani, G., Rehermann, M.I., Aldecosea, C., Trujillo-Cenóz, O., and Russo, R.E. (2018). Emergence of Serotonergic Neurons After Spinal Cord Injury in Turtles. Front Neural Circuits 12, 20. doi: 10.3389/fncir.2018.00020.

Faskowitz, J., Betzel, R.F., and Sporns, O. (2022). Edges in brain networks: Contributions to models of structure and function. Netw Neurosci 6(1), 1–28. doi: 10.1162/netn_a_00204.

Foote, S.L., and Morrison, J.H. (1984). Postnatal development of laminar innervation patterns by monoaminergic fibers in monkey (Macaca fascicularis) primary visual cortex. J Neurosci 4(11), 2667–2680.

Gagnon, D., and Parent, M. (2014). Distribution of VGLUT3 in highly collateralized axons from the rat dorsal raphe nucleus as revealed by single-neuron reconstructions. PLoS One 9(2), e87709. doi: 10.1371/journal.pone.0087709.

Gjorevski, N., Nikolaev, M., Brown, T.E., Mitrofanova, O., Brandenberg, N., DelRio, F.W., et al. (2022). Tissue geometry drives deterministic organoid patterning. Science 375(6576), eaaw9021. doi: 10.1126/science.aaw9021.

Goodfellow, I., Bengio, Y., and Courville, A. (2016). Deep Learning. Cambridge, MA: The MIT Press.

Greco, D., Volpicelli, F., Di Lieto, A., Leo, D., Perrone-Capano, C., Auvinen, P., et al. (2009). Comparison of gene expression profile in embryonic mesencephalon and neuronal primary cultures. PLoS One 4(3), e4977. doi: 10.1371/journal.pone.0004977.

Hawthorne, A.L., Wylie, C.J., Landmesser, L.T., Deneris, E.S., and Silver, J. (2010). Serotonergic neurons migrate radially through the neuroepithelium by dynamin-mediated somal translocation. J Neurosci 30(2), 420–430. doi: 10.1523/jneurosci.2333-09.2010.

Hendricks, T., Francis, N., Fyodorov, D., and Deneris, E.S. (1999). The ETS domain factor Pet-1 is an early and precise marker of central serotonin neurons and interacts with a conserved element in serotonergic genes. J Neurosci 19(23), 10348–10356.

Hornung, J.P. (2003). The human raphe nuclei and the serotonergic system. J Chem Neuroanat 26(4), 331–343. doi: 10.1016/j.jchemneu.2003.10.002.

Hrabetova, S., Cognet, L., Rusakov, D.A., and Nagerl, U.V. (2018). Unveiling the extracellular space of the brain: From super-resolved microstructure to in vivo function. J Neurosci 38(44), 9355–9363. doi: 10.1523/jneurosci.1664-18.2018.

Huang, C.X., Zhao, Y., Mao, J., Wang, Z., Xu, L., Cheng, J., et al. (2021). An injury-induced serotonergic neuron subpopulation contributes to axon regrowth and function restoration after spinal cord injury in zebrafish. Nat Commun 12(1), 7093. doi: 10.1038/s41467-021-27419-w.

Jacobs, B.L., and Azmitia, E.C. (1992). Structure and function of the brain serotonin system. Physiol Rev 72(1), 165–229. doi: 10.1152/physrev.1992.72.1.165.

Janušonis, S., and Detering, N. (2019). A stochastic approach to serotonergic fibers in mental disorders. Biochimie 161, 15–22. doi: 10.1016/j.biochi.2018.07.014.

Janušonis, S., Detering, N., Metzler, R., and Vojta, T. (2020). Serotonergic axons as fractional Brownian motion paths: Insights Into the self-organization of regional densities. Front Comput Neurosci 14, 56. doi: 10.3389/fncom.2020.00056.

Janušonis, S., Gluncic, V., and Rakic, P. (2004). Early serotonergic projections to Cajal-Retzius cells: relevance for cortical development. J Neurosci 24(7), 1652–1659. doi: 10.1523/jneurosci.4651-03.2004.

Janušonis, S., Mays, K.C., and Hingorani, M.T. (2019). Serotonergic axons as 3D-walks. ACS Chem Neurosci 10(7), 3064–3067. doi: 10.1021/acschemneuro.8b00667.

Jiang, X., Shen, S., Cadwell, C.R., Berens, P., Sinz, F., Ecker, A.S., et al. (2015). Principles of connectivity among morphologically defined cell types in adult neocortex. Science 350(6264), aac9462. doi: 10.1126/science.aac9462.

Jin, Y., Dougherty, S.E., Wood, K., Sun, L., Cudmore, R.H., Abdalla, A., et al. (2016). Regrowth of serotonin axons in the adult mouse brain following injury. Neuron 91(4), 748–762. doi: 10.1016/j.neuron.2016.07.024.

Johnson, M.D. (1994). Electrophysiological and histochemical properties of postnatal rat serotonergic neurons in dissociated cell culture. Neuroscience 63(3), 775–787. doi: 10.1016/0306-4522(94)90522-3.

Johnson, M.D., and Yee, A.G. (1995). Ultrastructure of electrophysiologically-characterized synapses formed by serotonergic raphe neurons in culture. Neuroscience 67(3), 609–623. doi: 10.1016/0306-4522(95)00010-g.

Kajstura, T.J., Dougherty, S.E., and Linden, D.J. (2018). Serotonin axons in the neocortex of the adult female mouse regrow after traumatic brain injury. J Neurosci Res 96(4), 512–526. doi: 10.1002/jnr.24059.

Katori, S., Hamada, S., Noguchi, Y., Fukuda, E., Yamamoto, T., Yamamoto, H., et al. (2009). Protocadherin-alpha family is required for serotonergic projections to appropriately innervate target brain areas. J Neurosci 29(29), 9137–9147. doi: 10.1523/jneurosci.5478-08.2009.

Katori, S., Noguchi-Katori, Y., Okayama, A., Kawamura, Y., Luo, W., Sakimura, K., et al. (2017). Protocadherin-alphaC2 is required for diffuse projections of serotonergic axons. Sci Rep 7(1), 15908. doi: 10.1038/s41598-017-16120-y.

Kiyasova, V., and Gaspar, P. (2011). Development of raphe serotonin neurons from specification to guidance. Eur J Neurosci 34(10), 1553–1562. doi: 10.1111/j.1460-9568.2011.07910.x.

Kosofsky, B.E., and Molliver, M.E. (1987). The serotoninergic innervation of cerebral cortex: different classes of axon terminals arise from dorsal and median raphe nuclei. Synapse 1(2), 153–168. doi: 10.1002/syn.890010204.

Labach, A., Salehinejad, H., and Valaee, S. (2019). Survey of dropout methods for deep neural networks. 1904.13310v2.

Lauder, J.M., Wallace, J.A., Krebs, H., Petrusz, P., and McCarthy, K. (1982). In vivo and in vitro development of serotonergic neurons. Brain Res Bull 9(1-6), 605–625. doi: 10.1016/0361-9230(82)90165-4.

Lautenschlager, M., Höltje, M., von Jagow, B., Veh, R.W., Harms, C., Bergk, A., et al. (2000). Serotonin uptake and release mechanisms in developing cultures of rat embryonic raphe neurons: age- and region-specific differences. Neuroscience 99(3), 519–527. doi: 10.1016/s0306-4522(00)00222-0.

Lavoie, B., and Parent, A. (1991). Serotoninergic innervation of the thalamus in the primate: an immunohistochemical study. J Comp Neurol 312(1), 1–18. doi: 10.1002/cne.903120102.

Lesch, K.P., and Waider, J. (2012). Serotonin in the modulation of neural plasticity and networks: implications for neurodevelopmental disorders. Neuron 76(1), 175–191. doi: 10.1016/j.neuron.2012.09.013.

Lidov, H.G., and Molliver, M.E. (1982). Immunohistochemical study of the development of serotonergic neurons in the rat CNS. Brain Res Bull 9(1-6), 559–604. doi: 10.1016/0361-9230(82)90164-2.

Linley, S.B., Hoover, W.B., and Vertes, R.P. (2013). Pattern of distribution of serotonergic fibers to the orbitomedial and insular cortex in the rat. J Chem Neuroanat 48-49, 29–45. doi: 10.1016/j.jchemneu.2012.12.006.

Liu, Y., and Nakamura, S. (2006). Stress-induced plasticity of monoamine axons. Front Biosci 11, 1794–1801. doi: 10.2741/1923.

Lu, J., Zhong, X., Liu, H., Hao, L., Huang, C.T., Sherafat, M.A., et al. (2016). Generation of serotonin neurons from human pluripotent stem cells. Nat Biotechnol 34(1), 89–94. doi: 10.1038/nbt.3435.

Lynn, C.W., and Bassett, D.S. (2019). The physics of brain network structure, function and control. Nat Rev Physics 1, 318–332.

Maddaloni, G., Bertero, A., Pratelli, M., Barsotti, N., Boonstra, A., Giorgi, A., et al. (2017). Development of serotonergic fibers in the post-natal mouse brain. Front Cell Neurosci 11, 202. doi: 10.3389/fncel.2017.00202.

Maddaloni, G., Migliarini, S., Napolitano, F., Giorgi, A., Nazzi, S., Biasci, D., et al. (2018). Serotonin depletion causes valproate-responsive manic-like condition and increased hippocampal neuroplasticity that are reversed by stress. Sci Rep 8(1), 11847. doi: 10.1038/s41598-018-30291-2.

Mai, J.K., and Ashwell, K.W.S. (2004). “Fetal Development of the Central Nervous System,” in The Human Nervous System, eds. G. Paxinos & J.K. Mai. Elsevier), 49–94.

Maia, G.H., Soares, J.I., Almeida, S.G., Leite, J.M., Baptista, H.X., Lukoyanova, A.N., et al. (2019). Altered serotonin innervation in the rat epileptic brain. Brain Res Bull 152, 95–106. doi: 10.1016/j.brainresbull.2019.07.009.

Mamounas, L.A., Blue, M.E., Siuciak, J.A., and Altar, C.A. (1995). Brain-derived neurotrophic factor promotes the survival and sprouting of serotonergic axons in rat brain. J Neurosci 15(12), 7929–7939.

Mercer, L.D., Higgins, G.C., Lau, C.L., Lawrence, A.J., and Beart, P.M. (2017). MDMA-induced neurotoxicity of serotonin neurons involves autophagy and rilmenidine is protective against its pathobiology. Neurochem Int 105, 80–90. doi: 10.1016/j.neuint.2017.01.010.

Migliarini, S., Pacini, G., Pelosi, B., Lunardi, G., and Pasqualetti, M. (2013). Lack of brain serotonin affects postnatal development and serotonergic neuronal circuitry formation. Mol Psychiatry 18(10), 1106–1118. doi: 10.1038/mp.2012.128.

Montgomery, T.R., Steinkellner, T., Sucic, S., Koban, F., Schüchner, S., Ogris, E., et al. (2014). Axonal targeting of the serotonin transporter in cultured rat dorsal raphe neurons is specified by SEC24C-dependent export from the endoplasmic reticulum. J Neurosci 34(18), 6344–6351. doi: 10.1523/jneurosci.2991-13.2014.

Morin, L.P., and Meyer-Bernstein, E.L. (1999). The ascending serotonergic system in the hamster: comparison with projections of the dorsal and median raphe nuclei. Neuroscience 91(1), 81–105. doi: 10.1016/s0306-4522(98)00585-5.

Moroz, L.L., Romanova, D.Y., and Kohn, A.B. (2021). Neural versus alternative integrative systems: molecular insights into origins of neurotransmitters. Philos Trans R Soc Lond B Biol Sci 376(1821), 20190762. doi: 10.1098/rstb.2019.0762.

Nazzi, S., Maddaloni, G., Pratelli, M., and Pasqualetti, M. (2019). Fluoxetine Induces Morphological Rearrangements of Serotonergic Fibers in the Hippocampus. ACS Chem Neurosci 10(7), 3218–3224. doi: 10.1021/acschemneuro.8b00655.

Nishi, M., Kawata, M., and Azmitia, E.C. (2000). Trophic interactions between brain-derived neurotrophic factor and s100beta on cultured serotonergic neurons. Brain Res 868(1), 113–118. doi: 10.1016/s0006-8993(00)02201-0.

Numasawa, Y., Hattori, T., Ishiai, S., Kobayashi, Z., Kamata, T., Kotera, M., et al. (2017). Depressive disorder may be associated with raphe nuclei lesions in patients with brainstem infarction. J Affect Disord 213, 191–198. doi: 10.1016/j.jad.2017.02.005.

Okaty, B.W., Commons, K.G., and Dymecki, S.M. (2019). Embracing diversity in the 5-HT neuronal system. Nat Rev Neurosci 20(7), 397–424. doi: 10.1038/s41583-019-0151-3.

Ren, J., Friedmann, D., Xiong, J., Liu, C.D., Ferguson, B.R., Weerakkody, T., et al. (2018). Anatomically defined and functionally distinct dorsal raphe serotonin sub-systems. Cell 175(2), 472-487.e420. doi: 10.1016/j.cell.2018.07.043.

Scheuch, K., Höltje, M., Budde, H., Lautenschlager, M., Heinz, A., Ahnert-Hilger, G., et al. (2010). Lithium modulates tryptophan hydroxylase 2 gene expression and serotonin release in primary cultures of serotonergic raphe neurons. Brain Res 1307, 14–21. doi: 10.1016/j.brainres.2009.10.027.

Senft, R.A., and Dymecki, S.M. (2021). Neuronal pericellular baskets: neurotransmitter convergence and regulation of network excitability. Trends Neurosci 44(11), 915–924. doi: 10.1016/j.tins.2021.08.006.

Šmít, D., Fouquet, C., Pincet, F., Zapotocky, M., and Trembleau, A. (2017). Axon tension regulates fasciculation/defasciculation through the control of axon shaft zippering. Elife 6. doi: 10.7554/eLife.19907.

Sporns, O., Tononi, G., and Kötter, R. (2005). The human connectome: A structural description of the human brain. PLoS Comput Biol 1(4), e42. doi: 10.1371/journal.pcbi.0010042.

Staal, R.G.W., Rayport, S., and Sulzer, D. (2007). Amperometric detection of dopamine exocytosis from synaptic terminals. In: Electrochemical Methods for Neuroscience, eds. A.C. Michael & L.M. Borland. CRC Press/Taylor & Francis.

Steinbusch, H.W. (1981). Distribution of serotonin-immunoreactivity in the central nervous system of the rat-cell bodies and terminals. Neuroscience 6(4), 557–618. doi: 10.1016/0306-4522(81)90146-9.

Stuesse, S.L., Cruce, W.L., and Northcutt, R.G. (1991). Localization of serotonin, tyrosine hydroxylase, and leu-enkephalin immunoreactive cells in the brainstem of the horn shark, Heterodontus francisci. J Comp Neurol 308(2), 277–292. doi: 10.1002/cne.903080211.

Sullivan, K.M., Ko, E., Kim, E.M., Ballance, W.C., Ito, J.D., Chalifoux, M., et al. (2022). Extracellular Microenvironmental Control for Organoid Assembly. Tissue Eng Part B Rev. doi: 10.1089/ten.TEB.2021.0186.

Sun, C., Qi, L., Cheng, Y., Zhao, Y., and Gu, C. (2022). Immediate induction of varicosities by transverse compression but not uniaxial stretch in axon mechanosensation. Acta Neuropathol Commun 10(1), 7. doi: 10.1186/s40478-022-01309-8.

Sundstrom, E., Kolare, S., Souverbie, F., Samuelsson, E.B., Pschera, H., Lunell, N.O., et al. (1993). Neurochemical differentiation of human bulbospinal monoaminergic neurons during the first trimester. Brain Res Dev Brain Res 75(1), 1–12. doi: 10.1016/0165-3806(93)90059-j.

Teuscher, C. (2022). Revisiting the edge of chaos: Again? Biosystems 218, 104693. doi: 10.1016/j.biosystems.2022.104693.

Vadodaria, K.C., Stern, S., Marchetto, M.C., and Gage, F.H. (2018). Serotonin in psychiatry: in vitro disease modeling using patient-derived neurons. Cell Tissue Res 371(1), 161–170. doi: 10.1007/s00441-017-2670-4.

van der Groen, O., Potok, W., Wenderoth, N., Edwards, G., Mattingley, J.B., and Edwards, D. (2022). Using noise for the better: The effects of transcranial random noise stimulation on the brain and behavior. Neurosci Biobehav Rev 138, 104702. doi: 10.1016/j.neubiorev.2022.104702.

Vertes, R.P. (1991). A PHA-L analysis of ascending projections of the dorsal raphe nucleus in the rat. J Comp Neurol 313(4), 643–668. doi: 10.1002/cne.903130409.

Vertes, R.P., Fortin, W.J., and Crane, A.M. (1999). Projections of the median raphe nucleus in the rat. J Comp Neurol 407(4), 555–582.

Voigt, T., and de Lima, A.D. (1991). Serotoninergic innervation of the ferret cerebral cortex. I. Adult pattern. J Comp Neurol 314(2), 403–414. doi: 10.1002/cne.903140214.

Vojta, T., Halladay, S., Skinner, S., Janušonis, S., Guggenberger, T., and Metzler, R. (2020). Reflected fractional Brownian motion in one and higher dimensions. Phys Rev E 102, 032108.

Vollenweider, F.X., and Kometer, M. (2010). The neurobiology of psychedelic drugs: implications for the treatment of mood disorders. Nat Rev Neurosci 11(9), 642–651. doi: 10.1038/nrn2884.

Voortman, L., and Johnston, R.J., Jr. (2022). Transcriptional repression in stochastic gene expression, patterning, and cell fate specification. Dev Biol 481, 129–138. doi: 10.1016/j.ydbio.2021.10.002.

Wada, A.H.O., and Vojta, T. (2018). Fractional Brownian motion with a reflecting wall. Phys Rev E 97(2-1), 020102. doi: 10.1103/PhysRevE.97.020102.

Wang, W., Pizzonia, J.H., and Richerson, G.B. (1998). Chemosensitivity of rat medullary raphe neurones in primary tissue culture. J Physiol 511 (Pt 2)(Pt 2), 433–450. doi: 10.1111/j.1469-7793.1998.433bh.x.

Way, B.M., Lacan, G., Fairbanks, L.A., and Melega, W.P. (2007). Architectonic distribution of the serotonin transporter within the orbitofrontal cortex of the vervet monkey. Neuroscience 148(4), 937–948. doi: 10.1016/j.neuroscience.2007.06.038.

Whitaker-Azmitia, P.M. (2001). Serotonin and brain development: role in human developmental diseases. Brain Res Bull 56(5), 479–485. doi: 10.1016/s0361-9230(01)00615-3.

Wisdom, K., and Chaudhuri, O. (2017). 3D Cell Culture in Interpenetrating Networks of Alginate and rBM Matrix. Methods Mol Biol 1612, 29–37. doi: 10.1007/978-1-4939-7021-6_3.

Yasufuku-Takano, J., Nakajima, S., and Nakajima, Y. (2008). Morphological and physiological properties of serotonergic neurons in dissociated cultures from the postnatal rat dorsal raphe nucleus. J Neurosci Methods 167(2), 258–267. doi: 10.1016/j.jneumeth.2007.08.018.

Yurchenko, I., Vensi Basso, J.M., Syrotenko, V.S., and Staii, C. (2019). Anomalous diffusion for neuronal growth on surfaces with controlled geometries. PLoS One 14(5), e0216181. doi: 10.1371/journal.pone.0216181.

